# Multi-epitope vaccine design using an immunoinformatics approach for 2019 novel coronavirus (SARS-CoV-2)

**DOI:** 10.1101/2020.03.03.962332

**Authors:** Ye Feng, Min Qiu, Liang Liu, Shengmei Zou, Yun Li, Kai Luo, Qianpeng Guo, Ning Han, Yingqiang Sun, Kui Wang, Xinlei Zhuang, Shanshan Zhang, Shuqing Chen, Fan Mo

## Abstract

A new coronavirus SARS-CoV-2 has caused over 9.2 million infection cases and 475758 deaths worldwide. Due to the rapid dissemination and the unavailability of specific therapy, there is a desperate need for vaccines to combat the epidemic of SARS-CoV-2. An in silico approach based on the available virus genome was applied to identify 19 high immunogenic B-cell epitopes and 499 human-leukocyte-antigen (HLA) restricted T-cell epitopes. Thirty multi-epitope peptide vaccines were designed by iNeo Suite, and manufactured by solid-phase synthesis. Docking analysis showed stable hydrogen bonds of epitopes with their corresponding HLA alleles. When four vaccine peptide candidates from the spike protein of SARS-CoV-2 were selected to immunize mice, a significantly larger amount of IgG in serum as well as an increase of CD19^+^ cells in ILNs was observed in peptide-immunized mice compared to the control mice. The ratio of IFN-γ-secreting lymphocytes in CD4^+^ or CD8^+^ cells in the peptides-immunized mice were higher than that in the control mice. There were also a larger number of IFN-γ-secreting T cells in spleen in the peptides-immunized mice. This study screened antigenic B-cell and T-cell epitopes in all encoded proteins of SARS-CoV-2, and further designed multi-epitope based peptide vaccine against viral structural proteins. The obtained vaccine peptides successfully elicited specific humoral and cellular immune responses in mice. Primate experiments and clinical trial are urgently required to validate the efficacy and safety of these vaccine peptides.

**Importance:** So far, a new coronavirus SARS-CoV-2 has caused over 9.2 million infection cases and 475758 deaths worldwide. Due to the rapid dissemination and the unavailability of specific therapy, there is a desperate need for vaccines to combat the epidemic of SARS-CoV-2. Different from the development approaches for traditional vaccines, the development of our peptide vaccine is faster and simpler. In this study, we performed an in silico approach to identify the antigenic B-cell epitopes and human-leukocyte-antigen (HLA) restricted T-cell epitopes, and designed a panel of multi-epitope peptide vaccines. The resulting SARS-CoV-2 multi-epitope peptide vaccine could elicit specific humoral and cellular immune responses in mice efficiently, displaying its great potential in our fight of COVID-19.

## Introduction

Since December 2019, a new type of coronavirus, SARS-CoV-2 (previously named 2019-nCoV by the World Health Organization), has caused an outbreak of viral lung infections in Wuhan City, Hubei Province, China, and later thrived over 30 countries worldwide (1-4). As of 24 June 2020, there have been over 9.2 million total confirmed cases and 475758 deaths of SARS-CoV-2 infection in the ongoing pandemic (5). Comparisons of the genome sequences of SARS-CoV-2 with other virus has shown 79.5% and 96% similarities at nucleotide level to SARS-CoV and bat coronaviruses, respectively (6), which suggested its probable origin in bats (7). The main clinical manifestations of SARS-CoV-2 patients are fever (≥38°C), dry cough, low or normal peripheral white blood cell count, and low lymphocyte count, known as novel coronavirus-infected pneumonia (NCIP) or coronavirus disease 2019 (COVID19) (8).

Currently, there is no approved therapeutics or vaccines available for the treatment of COVID-19 (9). Due to lack of anti-viral drugs or vaccines, control measures have been relying on the rapid detection and isolation of symptomatic cases (9). In this context, a safe and efficacious vaccine is urgently required. Traditional approaches for developing vaccines waste much time in isolating, inactivating and injecting the microorganisms (or portions of them) that cause disease. Fortunately, computation-based method enables us to start from analysis of viral genome, without the need to grow pathogens and therefore speeding up the entire process. Complete genome sequencing of SARS-CoV-2 has finished and paved the way for the vaccine development (9). The genome of SARS-CoV-2 encodes the spike protein, the membrane protein, the envelope protein, the nucleocapsid protein and a few replication and transcription-related enzymes. Given the lack of repairing mechanism of RNA virus replicase complex, mutations are prone to occur during virus replication. The 4% nucleotide difference of the virus isolated from Rhinolophus to that from human suggests that SARS-CoV-2 mutates rapidly to achieve the host conversion (10). Like SARS-CoV, SARS-CoV-2 uses its receptor binding domain (RBD) on the spike protein to bind to the host’s angiotensin-converting enzyme 2 (ACE2) (6, 9, 11, 12). The RBD of SARS-CoV-2 binds angiotensin-converting enzyme 2 (ACE2) with 10-20-fold higher affinity than does SARS-CoV (13). Consequently, the SARS-CoV-2 vaccine can be developed targeting the structural proteins, and in particular, the RBD region, following the strategy for the SARS-CoV vaccine development (11, 14-17).

An ideal vaccine may contain both B-cell epitopes and T-cell epitopes, with combination of which vaccine is able to either induce specific humoral or cellular immune against pathogens efficiently (18). Since the development of a peptide vaccine against the virus causing foot-and-mouth disease (19), the establishment of peptide synthesis method by Lerner et al. (20), along with the advent of a peptide vaccine design combining T-cell and B-cell epitopes, has accelerated the vaccine development. In the present study, we followed this in silico approach to identify the potential B-cell and T-cell epitope(s) from the spike, envelope and membrane proteins. Then we selected a few candidate vaccine peptides to immunize mice. As a result, these peptides successfully elicited specific humoral and cellular immune responses, showing their potentials in the real combat against SARS-CoV-2.

## Materials and Methods

### Data retrieval

The genome sequence of SARS-CoV-2 isolate Wuhan-Hu-1 was retrieved from the NCBI database under the accession number MN908947. Gene and protein sequences were acquired according to the annotation. In particular, the RBD region for the spike protein was referred to as the fragment from 347 to 520 amino acid (aa) (21).

### B-cell epitope prediction

The online tool in IEDB (Immune-Epitope-Database And Analysis-Resource) was used for the analysis of the conserved regions of the candidate epitopes (22). Prediction of linear B-cell epitopes was performed through Bepipred software (23). The antigenic sites were determined with Kolaskar method (24). The surface accessible epitopes were predicted by Emini tool (25).

### T-cell epitope prediction

The sequences of structural proteins were split into small fragments with a length of 9aa; their binding affinity with the 34 most prevalent HLA alleles was predicted using netMHCpan (26) and our in-house prediction software iNeo-Pred, respectively. iNeo-Pred was trained on a large immune-peptide dataset, and achieved a better performance in predicting binding affinity of epitopes to specific HLA alleles. Only the epitopes predicted by both tools were selected. Next, for each epitope, a HLA score was calculated based on the frequencies of binding HLA alleles in Chinese population, which will be used as metrics to select better candidates for downstream analysis.

### Vaccine peptide design

The vaccine peptides were designed by our in-house tool iNeo-Design. First, the selected B-cell epitopes and their adjacent T-cell epitopes were bridged to form candidate peptides with length no more than 30aa. Meanwhile, to facilitate the peptide synthesis, vaccine peptide sequences were optimized based on their hydrophobicity and acidity. To minimize the safety risk, peptides that contained toxicity potential, human homologous region (full-length matches and identity > 95%), or bioactive peptide were discarded.

Besides the vaccine peptides containing both B-cell epitopes and T-cell epitopes, iNeo-Design also utilized all predicted T-cell epitopes to generate T-cell epitopes-only vaccine peptides. For each vaccine candidate, the epitope counts and HLA score reflecting the population coverage were calculated. Vaccine candidates with the higher epitope counts and HLA score were considered to be preferable for the downstream analysis.

### Structural analysis

The online server swiss-model was used to predict the 3D protein structures of viral proteins and HLA molecules (27). The online server PEP_FOLD was used to predict T-cell epitopes’ structures (28). To display the interaction between T-cell epitopes and HLA molecules, T-cell epitope models were docked to HLA molecules using MDockPep (29). All predicted structures or models were decorated and displayed by the open source version of pymol program (https://github.com/schrodinger/pymol-open-source).

### Immuno-stimulation of B lymphocytes

The selected 4 peptides were synthesized with the solid phase synthesis method by GenScript Biotech Company (Nanjing, China), and were mixed at an equal concentration of 1mg/ml. The immunization experiment constituted three groups, each consisting of 12 6–8 week-old female C57mice. Mice were immunized subcutaneously with 100 μL of the following compounds: Group 1, 100 μg peptide mixture & 50 μl QuickAntibody-Mouse (Biodragen, Beijing, China); Group 2, 50 μl QuickAntibody-Mouse; and Group 3, 100 μl PBS as negative control. The immunization was performed once a week and repeated four times in total.

On the 14^th^, 21^st^, 28^th^ day after the 1^st^ immunization, retro-orbital blood was collected from 5 randomly selected mice in each group. The sera were tested for the presence of IgG by enzyme linked immunosorbent assay (ELISA). Briefly, ELISA plates were coated with 100 μL of S protein (2 μg/mL recombinant 2019-nCoV s-trimer protein; Novoprotein, Shanghai, China). The coated plates were incubated overnight at room temperature and washed with PBS containing 0.05% Tween 20 (PBS-T). Blocking buffer (1% nonfat milk in PBS) was added and incubated for 2 h at 37 °C. After washing with PBS-T, the diluted sera (1:1000 in PBS buffer containing 1% nonfat milk and 0.05% Tween-20) were added and incubated at 37 °C for 2 h. Next the plates were washed with PBS and again incubated at 37 °C for 2 h with horseradish peroxidase conjugated Goat Anti-mouse IgG antibody (1:5000; Genscript, Nanjing, China). Detection was carried out with O-Phenylene Diamine (OPD, 0.01%) substrate (Thermo Scientific, Waltham, US) for 30 min at 37 °C. Finally, the reaction was stopped using stop buffer (Solarbio, Beijing, China), and the absorbance was measured at 492 nm.

On the 28^th^ day after the 1^st^ immunization, 6 randomly selected mice were euthanized. The inguinal lymph nodes (ILNs) were harvested and processed into single cell suspensions. The cells were stained with Zombie Aqua (Biolegend, San Diego, US), APC-conjugated anti-mouse CD19 antibody (Biolegend, San Diego, US), PerCP/Cyanine5.5-conjugated anti-mouse CD95 (Fas) antibody (Biolegend, San Diego, US) and FITC-conjugated anti-mouse GL7 antibody (Biolegend, San Diego, US). The stained cells were resuspended in 500 μl PBS and subsequently processed by the Aria II flow cytometry instrument (BD, Franklin Lakes, US).

### Immuno-stimulation of T cells

The design of immunization experiment was similar to that for the B cells, but the injecting compounds were different: Group 1, 100 μg peptide mixture & 10 μg granulocyte-macrophage colony stimulating factor (GM-CSF; Novoprotein, Shanghai, China); Group2, 10 μg GM-CSF; Group 3, 100 μl PBS as negative control.

On the 14^th^ and 28^th^ day after the 1^st^ immunization, 3 randomly selected mice were euthanized, respectively. The relative proportions of T cells in the splenocytes and ILN lymphocytes were analyzed by the Aria II flow cytometry instrument (BD, Franklin Lakes, US). Briefly, spleen and ILN were harvested and processed into single cell suspensions. Splenocytes (1×10^6^/well) and ILN lymphocyte (1×10^6^/well) were cultured overnight with the peptide mixture (5μg/ml) or in RPMI-1640 alone (negative control). Cells were stained with Zombie Aqua (Biolegend, San Diego, US), PerCP/Cyanine5.5-conjugated anti-mouse CD8a antibody (Biolegend, San Diego, US), PE/Cy7-conjugated anti-mouse CD4 antibody (Biolegend, San Diego, US) and APC-conjugated anti-mouse IFN-γ antibody (Biolegend, San Diego, US). The stained cells were resuspended in 500μl PBS for flow cytometry analysis.

The IFN-γ-secreting T lymphocytes were also quantified on 6 randomly selected mice using an ELISPOT kit (Dakewe, Shenzhen, China). Briefly, 100 μL of PBS were added to 96-well plates pre-coated with an anti-IFN-γ mAb. 2×10^5^/well cells were incubated in duplicate with 5μg/mL of peptide or medium alone (negative control) for 16 h in a 37 °C humidified incubator with 5% CO_2_. Splenocytes stimulated with phorbol myristate acetate (PMA) served as positive control. After removing cells and washing with buffer (PBS with 0.1% Tween 20), 1:100 diluted biotinylated anti-IFN-γ were added and incubated for 90 min at 37°C. After each incubation step, the plates were washed three times with buffer. Next, after 1 h of incubation with streptavidin-alkaline phosphatase conjugate (1/5000 in PBS-0.1% Tween 20), the plates were developed with a solution of 5-bromo-4-chloro-3-indolylphosphate (BCIP)-nitroblue tetrazolium until red spots appeared. Tap water was used to stop the reaction, and the plates were dried in air overnight. Individual spots were counted under a CTL-ImmunoSpot^®^S6 FluoroSpot (Cellular Technology, Kennesaw, USA).

### Statistical analysis

Comparisons were analyzed by one-way analysis of variance (ANOVA). P value less than 0.05 was considered significant.

## Results

### Prediction of B-cell epitopes

During the immune response against viral infection, B-cell takes in viral epitopes to recognize viruses and activates defense responses. Recognition of B-cell epitopes depends on antigenicity, accessibility of surface and predictions of linear epitope (30). A total of 61 B-cell epitopes were predicted, which seemed preferentially located within certain regions of each gene (Figure 1; Figure 2; Table S1). Only 19 epitopes were exposed on the surface of the virion and had a high antigenicity score, indicating their potentials in initiating immune response. Therefore, they were considered to be promising vaccine candidates against B-cells. Among the 19 epitopes, 17 were longer than 14 residues and located in the spike protein that contained RBD and functioned in host cell binding (Table 1). The average Emini score for the 19 epitopes was 2.744, and the average for Kolaskar (antigenicity) score was 1.015. Two epitopes were located within the RBD region, while the one with the highest Kolaskar score (1.059), 1052-FPQSAPH-1058, was located at position 1052aa of the spike protein.

**Table 1.**
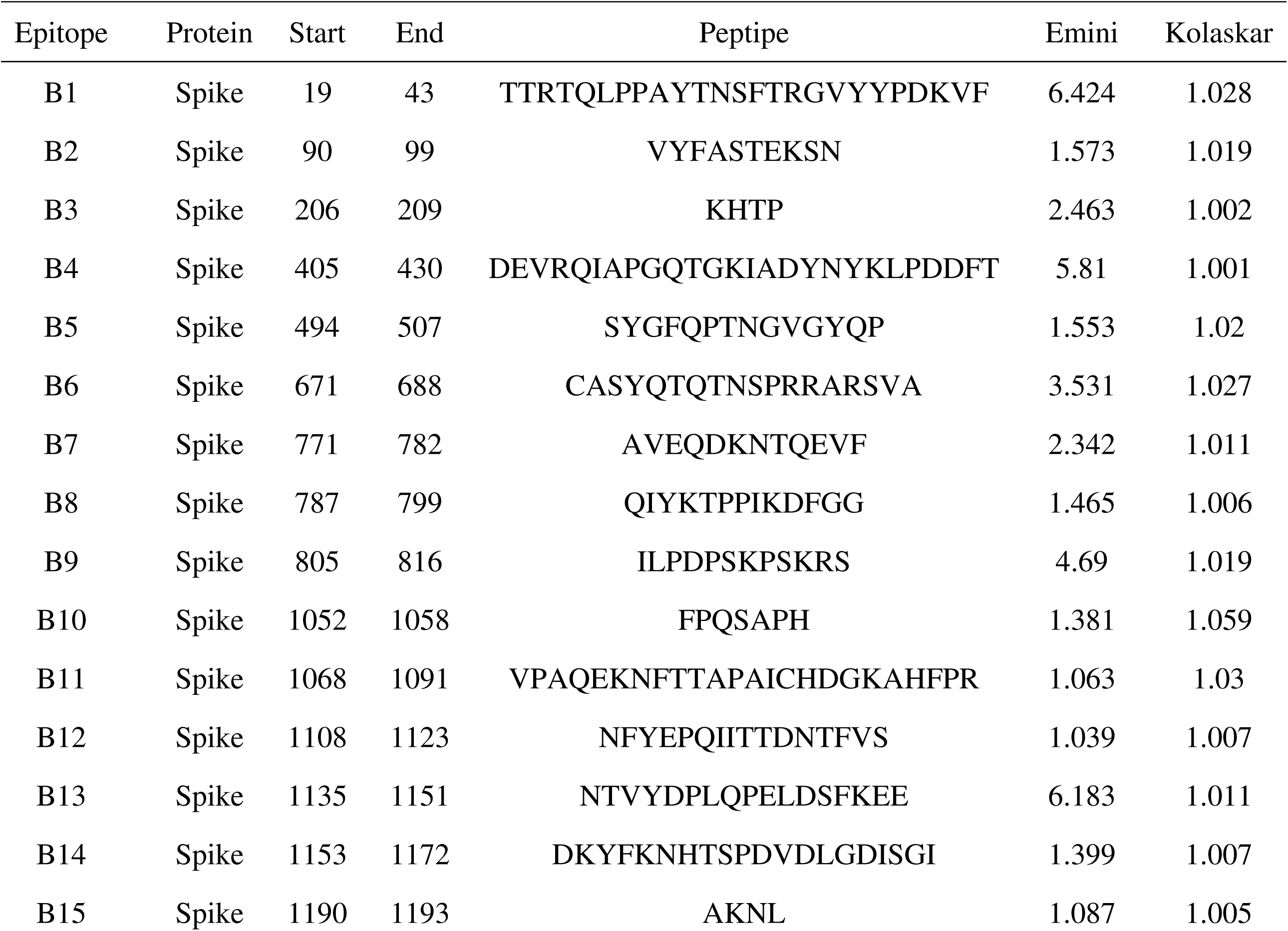
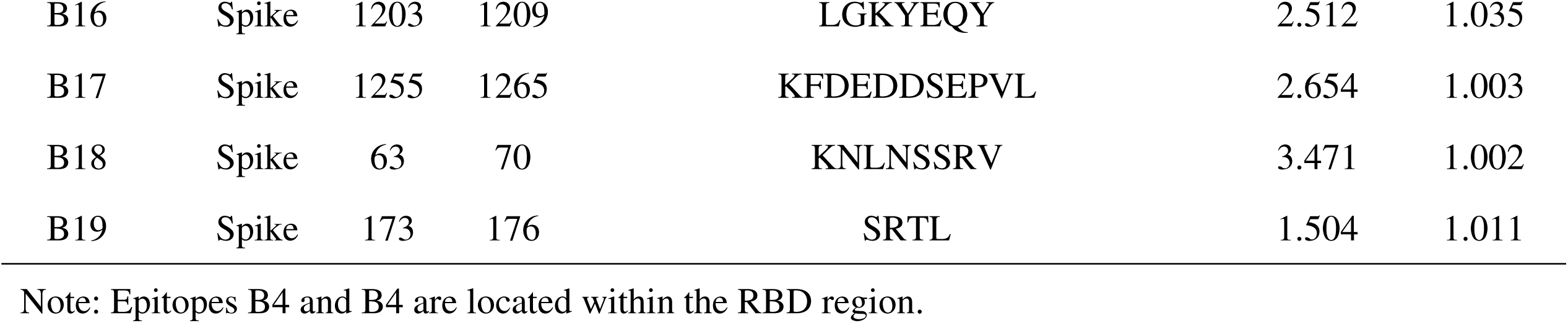
B-cell epitope candidates.

**Fig. 1.**
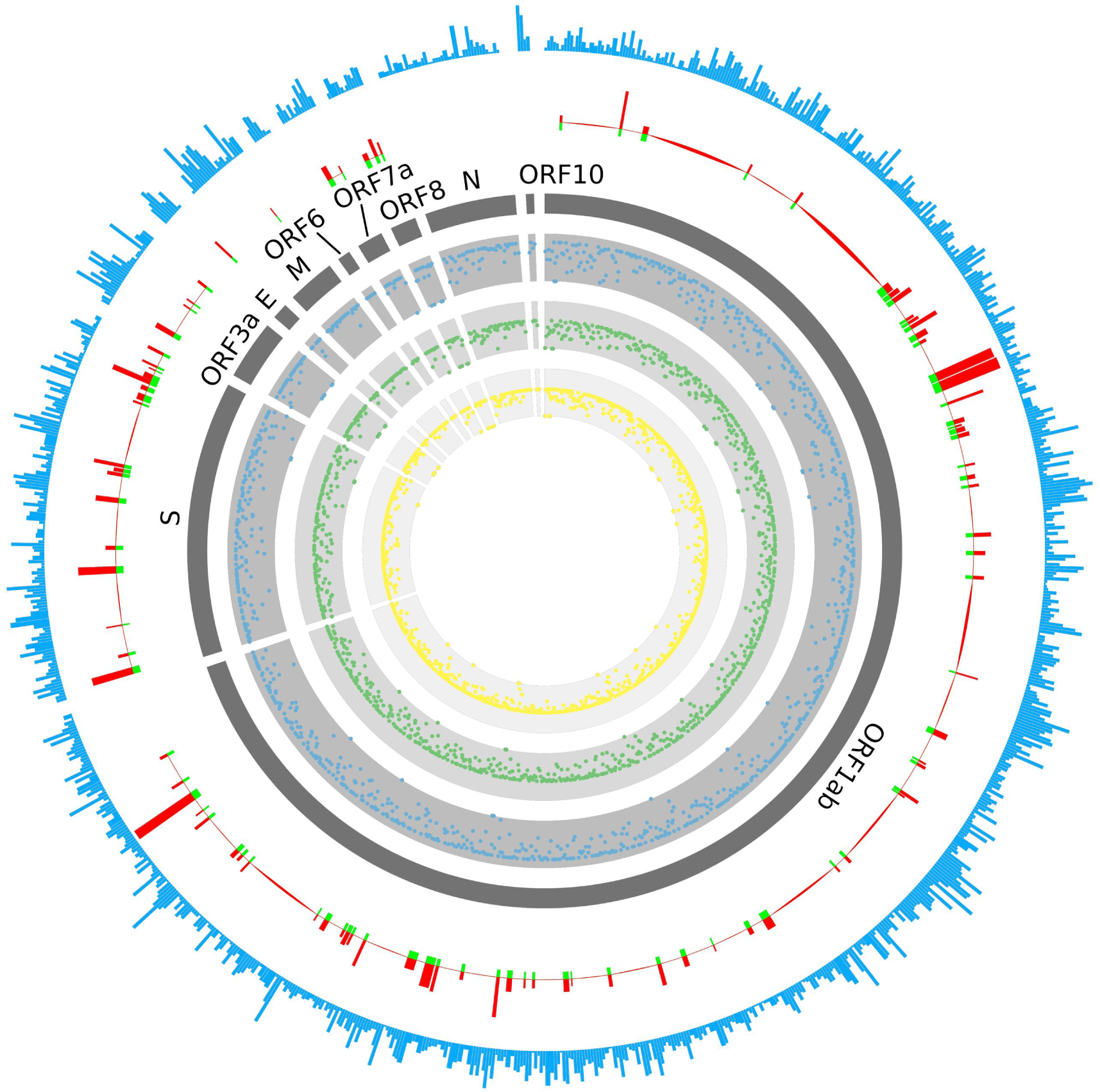
Distribution of B-cell and T-cell epitopes. The outermost circle (light blue) stands for the T-cell epitope count. The 2rd outer circle stands for Emini (in red) and Kolaskar (in green) score used to evaluate the B-cell epitopes. The 3rd circle marked the name of the viral proteins. The 4th-6th circles stands for HLA-A (in blue), HLA-B (in green), and HLA-C (in yellow) scores; the points closer to the center indicates a lower score.

**Fig. 2.**
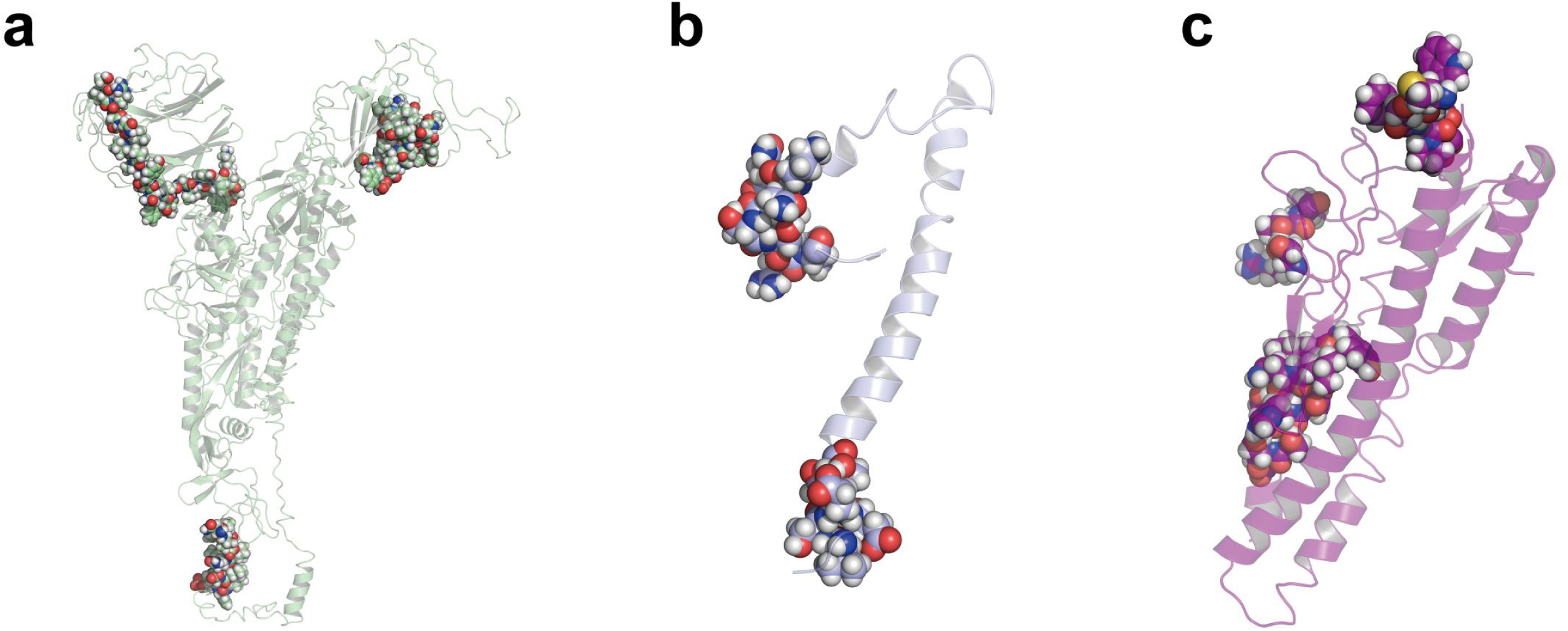
Locations of the recognized B cell epitopes on the viral spike protein (a), envelop protein (b) and membrane protein (c). The transparent cartoon models display the predicted 3D structure; the colorful balls marks the position of the recognized epitopes.

### Prediction of T-cell epitopes

The immune response of T-cell is considered as a long lasting response compared to B-cell where the antigen might easily escape the antibody memory response (31). Moreover, the CD8+ T and CD4+ T-cell responses play a major role in antiviral immunity. It is therefore important to design vaccines that can induce T-cell’s immune response (32). A total of 499 T-cell epitopes were predicted on the spike protein (378 epitopes), the membrane protein (90 epitopes) and the envelop protein (31 epitopes); 48 of the 378 epitopes for the spike protein were located in the RBD region (Figure 1; Table 2; Table S2). There is no preference in certain genes or regions for T-cell epitope generation; no biased distribution of T-cell epitopes among HLA types was observed either. Among all T-cell epitopes, the epitope 869-MIAQYTSAL-877 in the spike protein was predicted to be able to bind to 17 HLA alleles. Most of the HLA alleles included in the present study were covered by these vaccine candidates, which suggested a wide population coverage.

**Table 2.** Distribution of T-cell epitopes among three structural proteins.

| **Protein** | **Count of T-cell**<br>**Epitope** | **No. of epitope**<br>**per residue** | **Epitope overage** | **HLA**<br>**Types**<br>**Count** |
| --- | --- | --- | --- | --- |
| Spike | 378 | 0.297 | 93.01% | 33 |
| Membrane | 90 | 0.405 | 96.00% | 31 |
| Envelope | 31 | 0.413 | 94.14% | 32 |

In terms of the distribution of the predicted epitopes against different HLA haplotypes, no significant differences were observed among different HLA haplotypes (Table S3). There were 287, 208 and 195 epitopes predicted to be able to bind to HLA-A, HLA-B and HLA-C haplotypes, respectively. For the most popular five HLA types (HLA-A*11:01, HLA-A*24:02, HLA-C*07:02, HLA-A*02:01 and HLA-B*46:01), the counts for epitopes with binding affinity were 51, 49, 115, 48 and 58.

### Multi-epitope vaccine design

Based on the 19 B-cell epitopes and their 121 adjacent T-cell epitopes, 17 candidate vaccine peptides that contained both B-cell and T-cell epitopes were generated by our in-house software iNeo-Design. Most of the 17 candidate vaccine peptides contained one B-cell epitopes, except for AVEQDKNTQEVFAQVKQIYKTPPIKDFGG, which involved two B-cell epitopes and eight T-cell epitopes, and AKNLNESLIDLQELGKYEQYIKWPWYIWKK, which contained two B-cell epitopes and 6 T-cell epitopes. By comparison, the vaccine peptide FKNLREFVFKNIDGYFKIYSKHTPINLV had the largest count of T-cell epitopes, whereas the vaccine peptide SYGFQPTNGVGYQPYRVVVLSFELLHAPAT showed the highest HLA score, indicating their wide population coverage and promising efficacy.

In addition to the vaccine candidates involved both B-cell and T-cell epitopes, we also analyzed the entire 499 core T-cell epitopes to generate another 102 vaccine peptides containing T-cell epitopes only. Based on both the epitope counts and HLA score, we eventually selected 13 T-cell epitopes-only vaccine peptides.

Taken together, a total of 30 peptide vaccine candidates were designed (Table 3). 26 of them were from the spike protein, two from the membrane protein and two from the envelope protein. Five peptides were located in the RBD region, indicating they were likely to induce the production of neutralizing antibody. The 30 vaccine peptides covered all structural proteins that may induce immune response against SARS-CoV-2 in theory; and the multi-peptide strategy we applied would better fit the genetic variability of the human immune system and reduce the risk of pathogen’s escape through mutation (33).

### Interaction of predicted peptides with HLA alleles

To further inspect the binding stability of T-cell epitopes against HLA alleles, the T-cell epitopes involved in the above designed vaccine peptides were selected to conduct an interaction analysis. Figure 3 illustrated the docking results against the most popular HLA types for the two epitopes from vaccines peptide 25 and 27 (Table 3; Table 4), which showed relatively higher HLA score. The MDockPep scores were between -148 ∼ -136, indicating that the predicted crystal structures were stable. All epitopes were docked inside the catalytic pocket of the receptor protein. In particular, the epitope 1220-FIAGLIAIV-1228 from the spike protein possessed 2-5 stable hydrogen bonds with the HLA alleles; the epitope 4-FVSEETGTL-12 from the envelop protein possessed 4∼5 stable hydrogen bonds (Table 4). Taken together, the epitopes included in our vaccine peptides can interact with the given HLA alleles by in silico prediction.

**Table 3.**
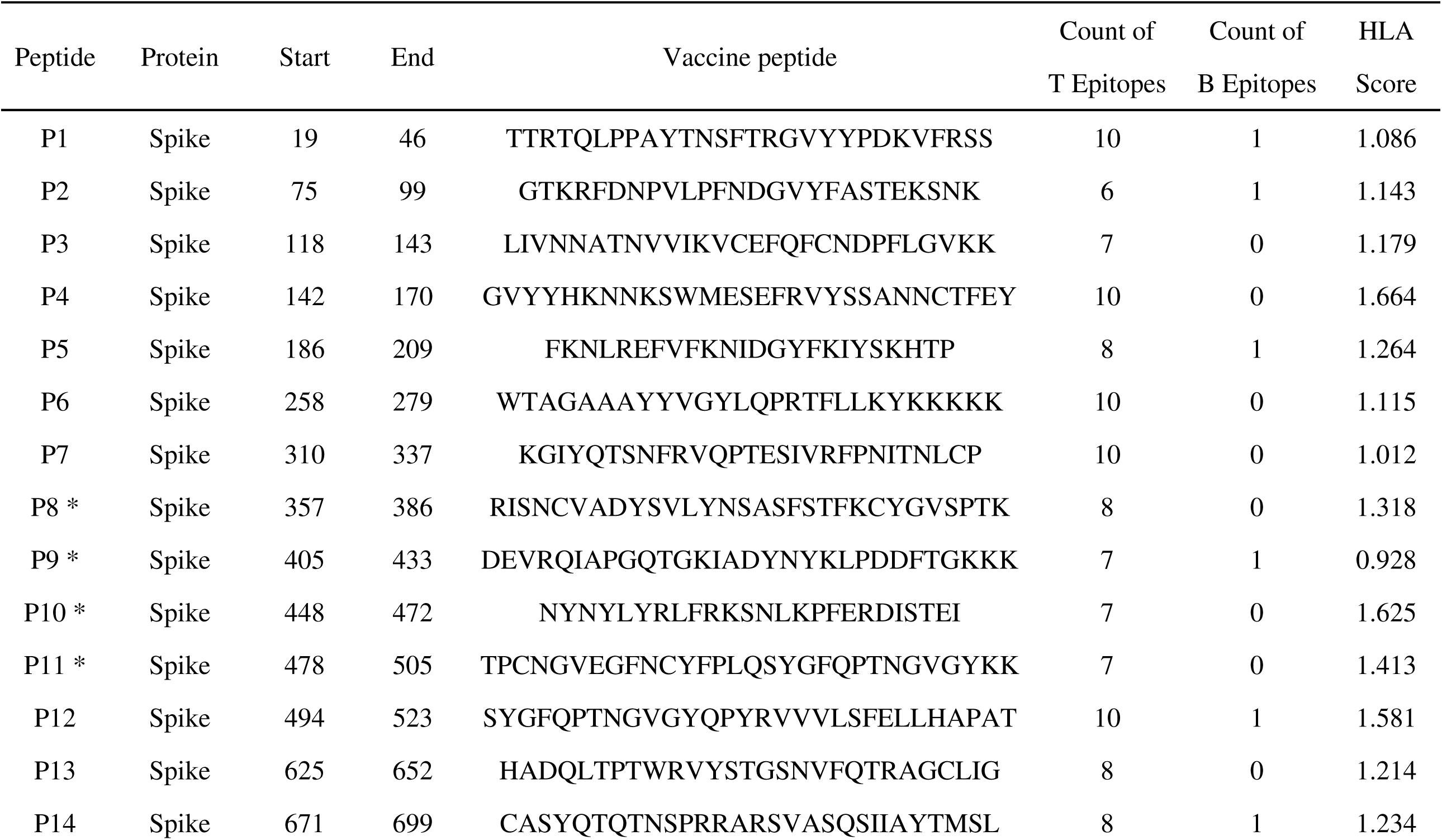
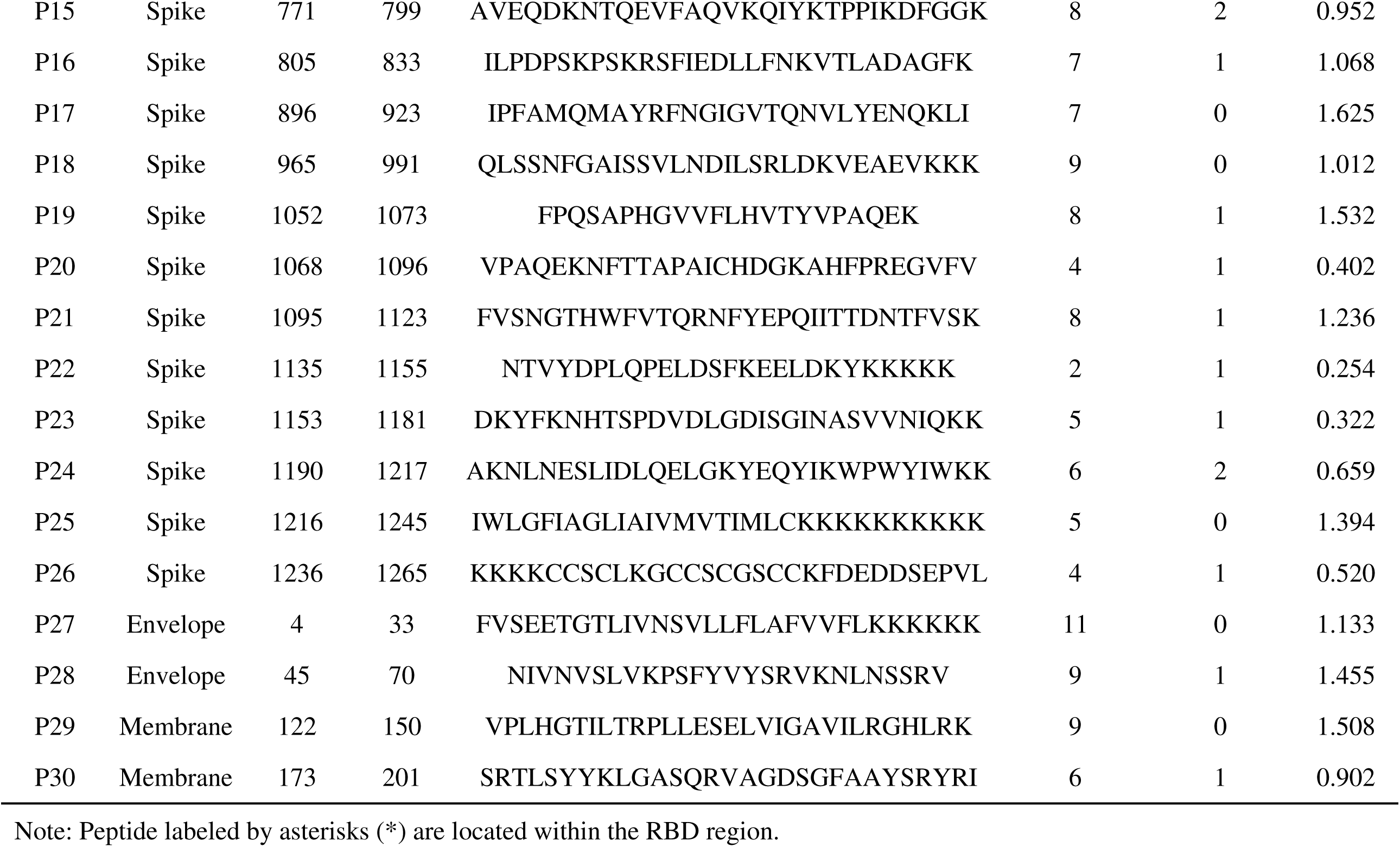
Candidate vaccine peptides.

**Table 4.**
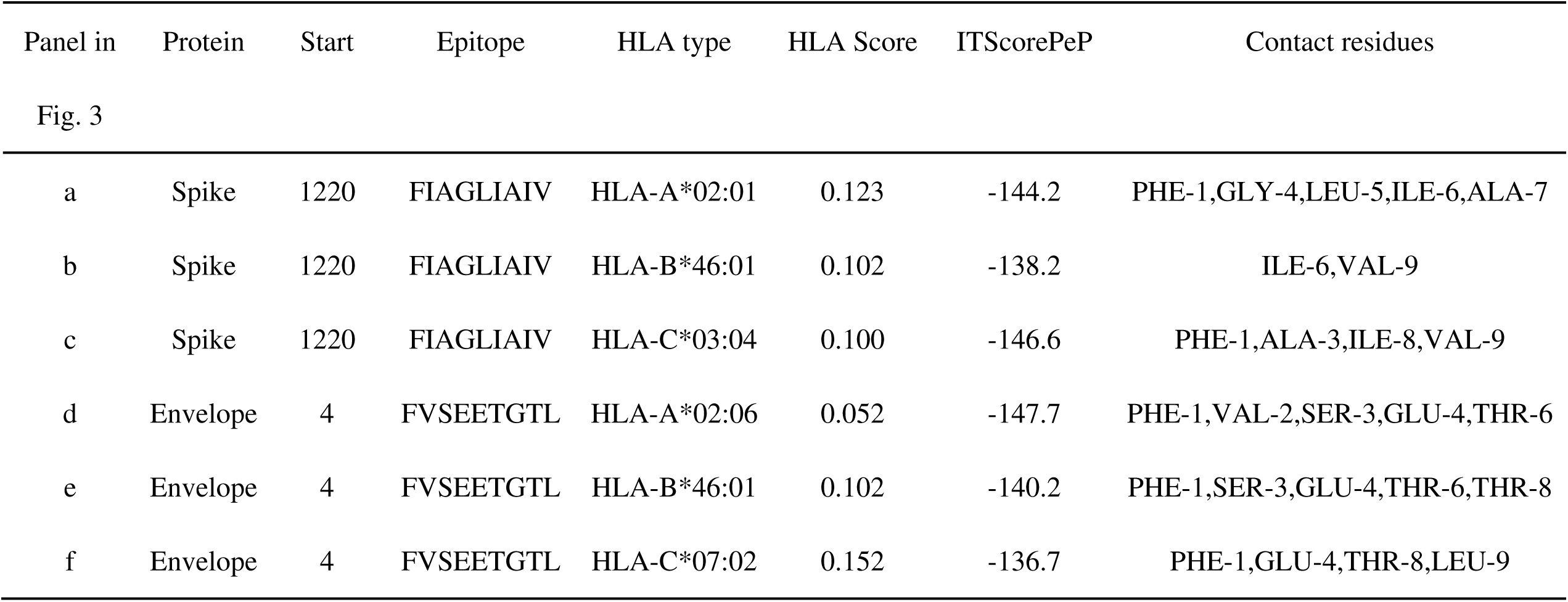
Docking results for T-cell epitope P25 and P27 against three HLA types.

**Fig. 3.**
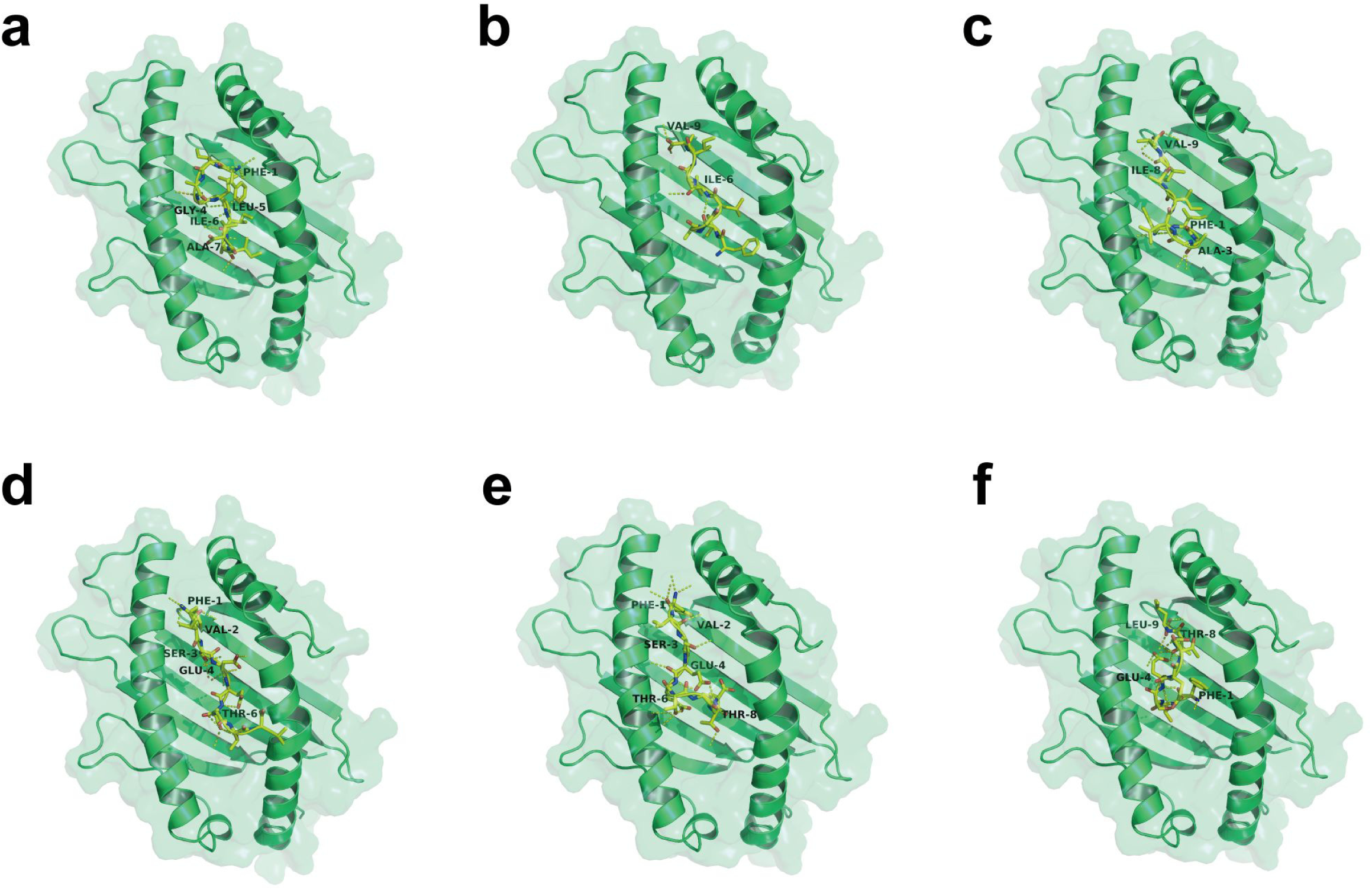
Interaction between the predicted peptides (by yellow sticks) and different HLA alleles (by green cartoons). Amino acids were labeled adjacent to the contact sites. Table 3 displays the detailed docking information.

### Humoral immune responses to SARS-CoV-2 S protein

Based on the above immunoinformatics analysis, 4 designed vaccine peptides, namely P9, P12, P14 and P15, were chosen as the candidates for the downstream validation experiments because of their relatively higher counts of B-cell and T-cell epitopes and the higher frequencies of their epitopes’ corresponding HLA alleles (Table 3). We immunized mice by subcutaneous injection of the mixture of these synthesized peptides plus QuickAntibody (an adjuvant for stimulating B cells). Mice injected with QuickAntibody only or PBS were studied as controls. The immunization was performed once a week and repeated four times in total.

To evaluate whether these peptides induce B cells to produce specific antibody against the S protein, an ELISA assay was conducted to detect IgG in the sera of mice. Fourteen days after the 1^st^ immunization, the amount of IgG showed little difference between the peptide-treated mice and the controls (Figure 4a), suggesting that two weeks were not long enough to elicit humoral immune response. Since the 21^st^ day after the 1^st^ immunization, however, the expression of IgG had risen to the plateau in the peptide-treated mice and was remarkably higher than that in the control mice (p<0.05; Figure 4a). Germinal centers (GCs) are the main sites for the production of high-affinity, long-lived plasma cells and memory B cells. On the 28^th^ day after the 1^st^ immunization, we collected ILN and stained cells with antibody of GL7 and FAS. Flow cytometry showed that there were much more B cells activated in the peptides-treated mice than in the mice injected with adjuvant only (Figure 4b); and the numbers of rapidly proliferated B cells (CD19^+^/FAS^+^/GL7^+^) from GC in ILNs of the peptides-treated mice were significantly higher than that in the control groups, demonstrating that there were increased GCs induced by peptide vaccines (Figure 4c-d). In future, a viral neutralization study is further required for demonstrating that the designed peptide vaccines can efficiently activate specific humoral immune responses to the S protein of SARS-CoV-2.

**Fig. 4.**
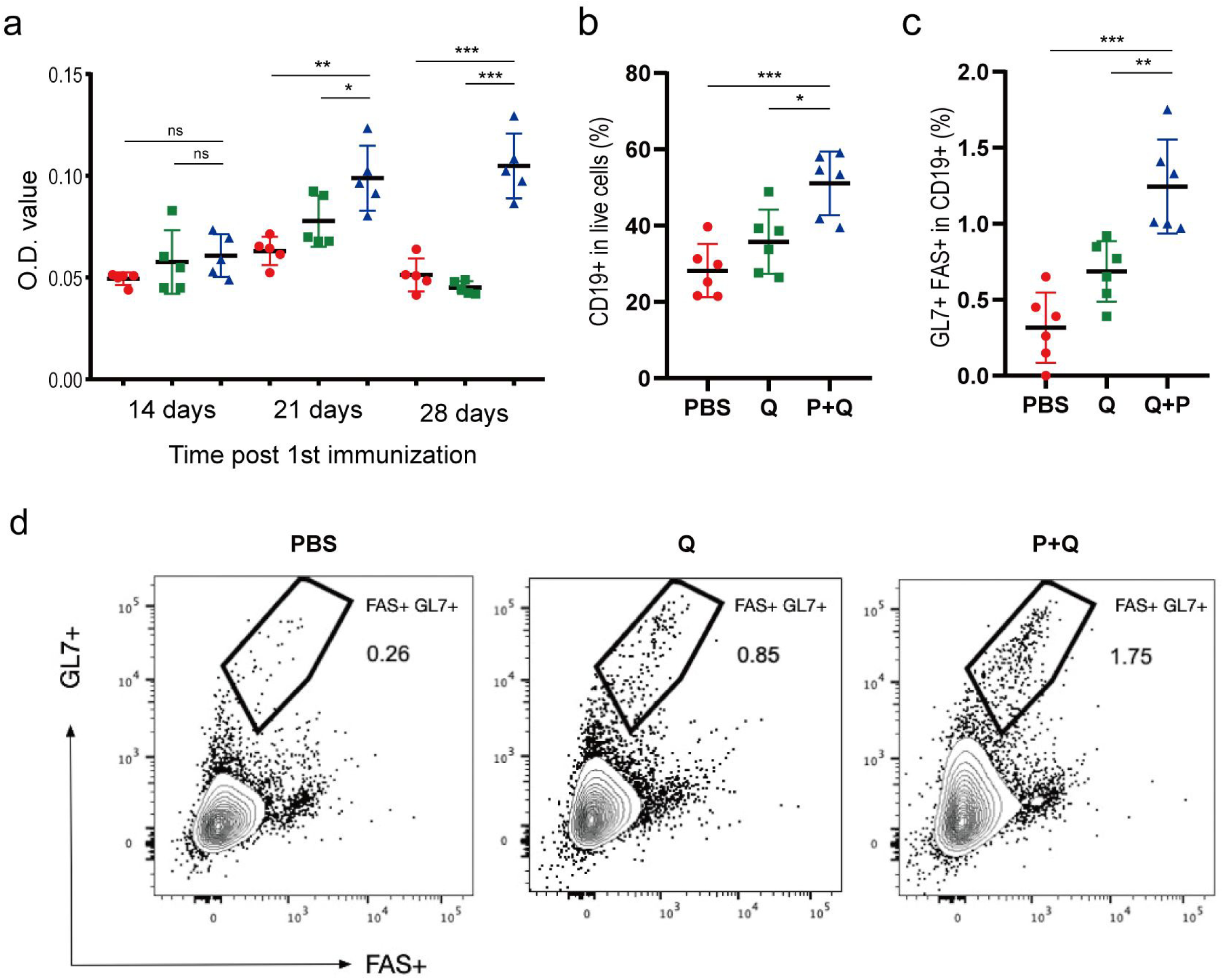
Humoral immune responses to SARS-CoV-2 S protein. (a) Comparison of humoral response among groups of mice injected with PBS (marked in red), QuickAntibody (in green) and Peptide + QuickAntibody (in blue), respectively. The level of IgG was measured by ELISA. (c) The percentage of B cells (CD19^+^ cells) in live cells. (c) The percentage of GC cells (FAS^+^/GL7^+^ cells) in CD19^+^ cells. (d) Flow cytometry showing the larger number of FAS^+^/GL7^+^ cells in the peptides-treated mice. PBS, Q, P+Q represent mice injecting with PBS, mice with QuickAntibody, and mice with peptide vaccines plus QuickAntibody, respectively. *, p<0.05; **, p<0.01; ***, p<0.001; ns, not significant.

### Cellular immune responses to SARS-CoV-2 S protein

In parallel, we also immunized mice with peptides plus GM-CSF, an adjuvant that induced the development of monocytes, neutrophils and dendritic cells. Mice injected with GM-CSF only or PBS were used as controls. On both the 14^th^ and the 28^th^ day after the 1^st^ immunization, the ILNs were collected. We found that the ratios of IFN-γ-secreting cells to both CD4^+^ and CD8^+^ T cells in the peptides-treated mice were significantly higher than that in the control groups (Figure 5), suggesting the activation of T cells by the peptide vaccines. Notably, the ratio of IFN-γ-secreting cells seemed to reach its plateau on the 14^th^ day, since no further significant increase of this ratio was observed on the 28^th^ day. It was likely that, in the absence of the virus, the repeated T cell stimulation led to depletion or transfer of T cells in ILNs; accordingly, the ratio of IFN-γ-secreting cells in the ILNs kept relatively stable afterwards.

**Figure 5.**
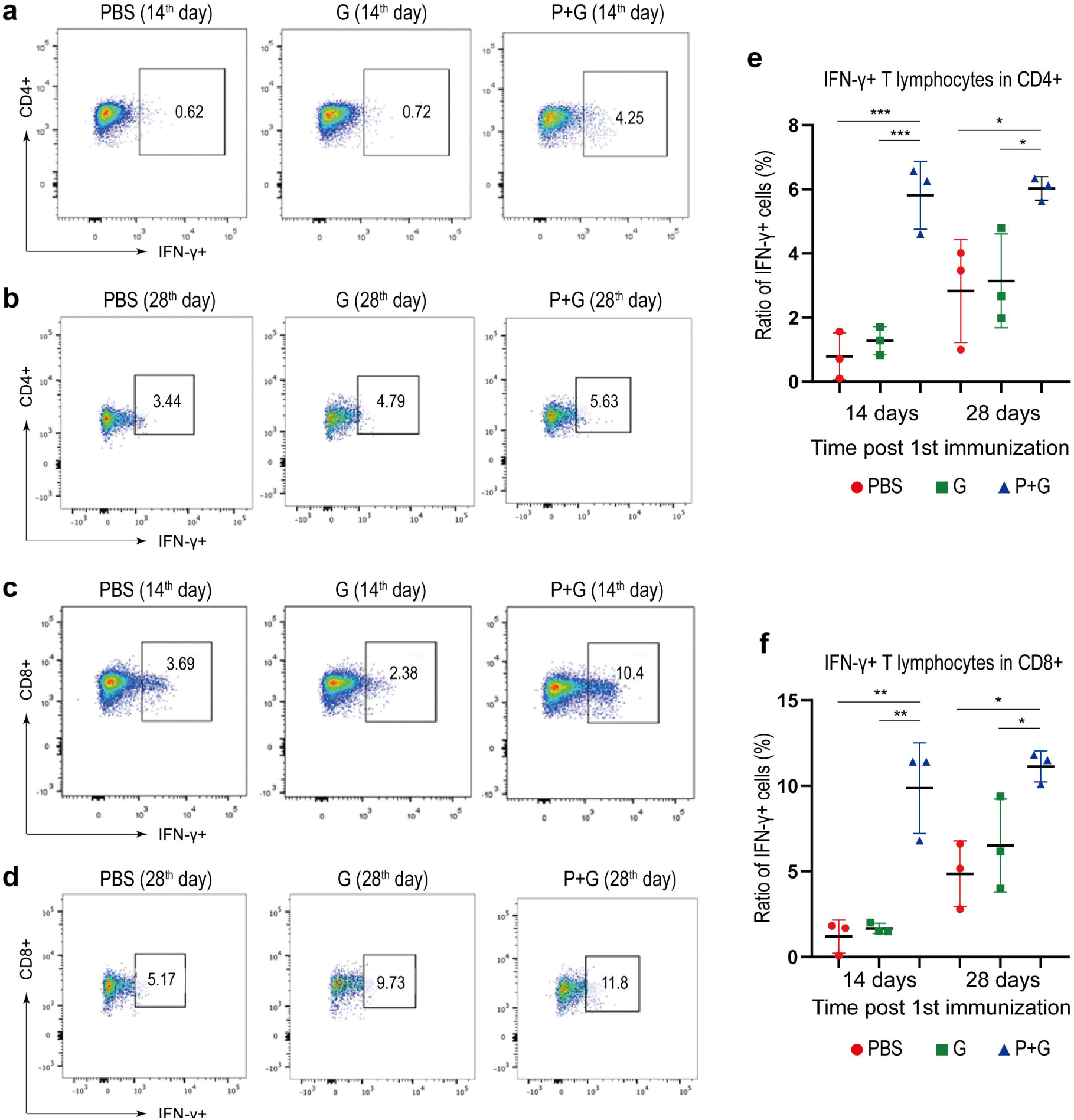
Specific T cell activation by SARS-CoV-2 S protein. (a) and (b) The IFN-γ-secreting T lymphocytes in CD4^+^ cells on the 14^th^ and the 28^th^ day after the 1^st^ immunization, respectively. (c) and (d) The IFN-γ-secreting T lymphocytes in CD8^+^ cells on the 14^th^ and the 28^th^ day after the 1^st^ immunization, respectively. (e) and (f) The percentage of IFN-γ-secreting cells in CD4^+^ cells and CD8^+^ cells, respectively. PBS, G, P+G represent mice injected with PBS, GM-CSF, and peptide vaccines plus GM-CSF, respectively. *, p<0.05; **, p<0.01; ***, p<0.001.

**Figure 6.**
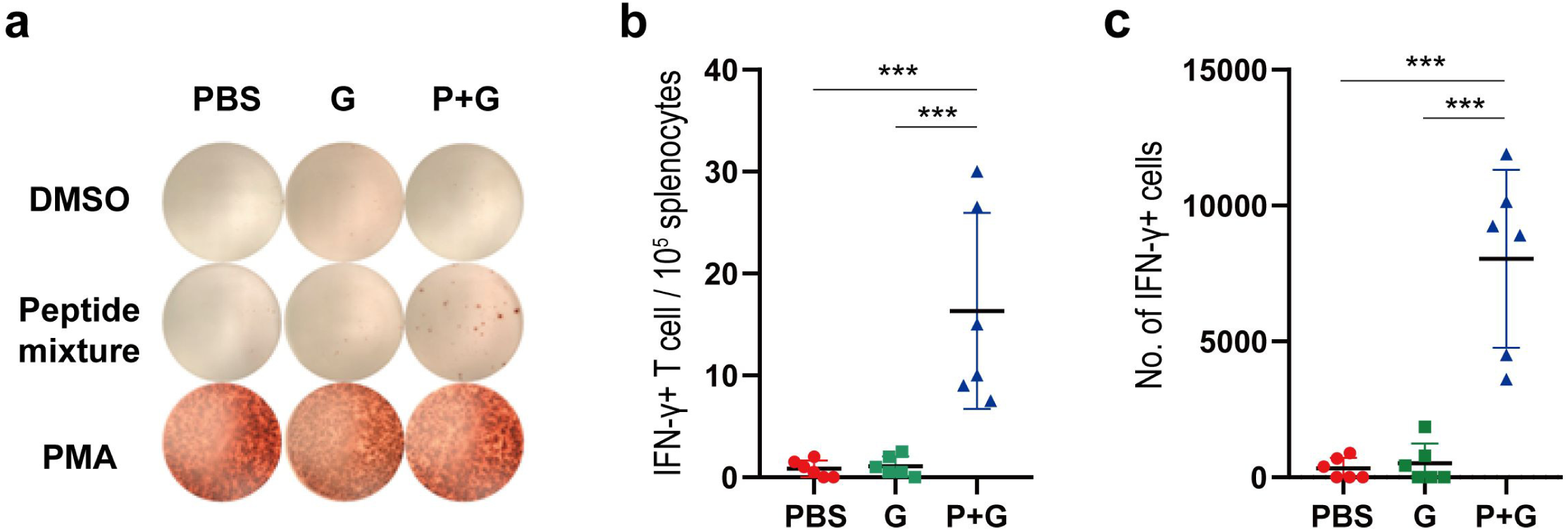
Quantification of the IFN-γ-secreting lymphocytes in mice spleen by ELISPOT. (a), Responses of splenocyte to DMSO (negative control), PMA (positive control) and the peptide mixture. (b) The number of IFN-γ-secreting cells per 100,000 splenocytes. (c) Total number of IFN-γ-secreting cells in spleen. PBS, G, P+G represent mice injected with PBS, GM-CSF, and peptide vaccines plus GM-CSF, respectively. ***, p<0.001.

We also quantified the peptide specific lymphocytes in mice spleen on the 28^th^ day using the enzyme-linked immunospot (ELISPOT) assay of IFN-γ secreting. In terms of either the ratio of IFN-γ-secreting lymphocytes in splenocytes or the total number of IFN-γ-secreting lymphocytes in spleen, mice immunized with peptide vaccines had significantly higher ratios than that in the control groups. This finding was overall consistent with the flow cytometry results of ILN cells, suggesting that the lymphocytes were activated and might recirculate to gather in the spleen after 4-week vaccination.

## Discussion

Since the outbreak of COVID-19, numerous researchers have contributed in the development of safe and effective vaccines for COVID-19 to increase the chances of success. Meanwhile, World Health Organization (WHO) has encouraged scientists to test all candidate vaccines until they fail. Currently, the developing candidate vaccines include inactivated virus vaccine, recombinant protein vaccine, viral vector vaccine, peptide vaccine, DNA and RNA vaccines.

In comparison, the involvement of the optimization of cell and virus culture process increase the complexity and thus preparation time of inactivated virus vaccine and viral vector vaccine. Also, for the development of inactivated virus vaccine, besides Good Manufacturing Practices (GMP) system, extremely high manufacture standard is required to avoid medical accidents due to the failure of complete inactivation of virus toxicity. Differently, the development of our peptide vaccine is much simpler, including only two major steps: sequence design through reverse vaccinology approach, and peptide synthesis.

Until now, most recombinant protein vaccine candidates have focused on spike protein, which although can provide good safety, is probably unable to stimulate strong T cell immune response (34). Therefore, effective adjuvants are usually added when use this kind of vaccines. Similar to recombinant protein vaccines, most viral vector vaccine candidates are based on the expression of spike protein (35). Although the application of viral vector could enhance the vaccine’s delivery efficiency, the body’s immune response to vector itself might interfere with the immune response to target epitopes, and thus compromise the efficacy of this kind of vaccines. Unlike the vaccine candidates discussed above, our peptide vaccine candidate is composed of 30 peptides from not only spike protein, but also membrane protein and envelope protein of SARS-CoV-2, containing both B-cell epitopes and T-cell epitopes to induce specific humoral and cellular immune response against SARS-CoV-2 more efficiently.

On the other hand, although promising in preclinical or Phase I and II clinical trials, no mRNA vaccine has been formally approved by FDA (36). In other words, more clinical trials should be conducted to prove the efficacy of mRNA vaccines. Moreover, the batch production, manufacturing process, and industrial expansion of mRNA vaccines are still under exploration. For DNA vaccines, besides the same concerns for mRNA vaccines, the possibility of the integration of foreign DNA sequence to human genome has brought uncertainty in their development. Different from these nucleic acid vaccines, peptide vaccine has relatively more mature manufacturing process. And the successful launch of previous peptide vaccines has already demonstrated the safety and efficacy of peptide-based vaccines.

In this study, our SARS-CoV-2 multi-epitope peptide vaccine could elicit specific humoral and cellular immune responses in mice, displaying its great potential in the fight of COVID-19. In future, more experiments will be conducted to validate the efficacy of our SARS-CoV-2 peptide vaccine.

## Authors’ contributions

SC and FM conceived and designed the project. YF, FM, MQ, SZ, KL, YS, KW, XZ and SZ analyzed the data. LL and QG conducted in-vivo validation. YF, YL and NH wrote the initial draft. All authors revised and approved the final manuscript.

## Acknowledgement

The authors would like to thank Dr. Jing Guo for the suggestion on the in-vivo experiments.

## Ethics approval and consent to participate

Ethical approval for the animal experiments was obtained from Zhejiang Chinese Medical University Laboratory Animal Research Center.

## Competing interests

The authors declare that they have no competing interests.

## Notes

### Competing Interest Statement

The authors have declared no competing interest.

### Summary of Updates

A few candidate vaccine peptides were selected to immunize mice. As a result, these peptides successfully elicited specific humoral and cellular immune responses, showing their potentials in the real combat against SARS-CoV-2. The relevant experiments and results were added in this manuscript.

