## Supplemental material for "Multi-epitope vaccine design using an immunoinformatics approach for 2019 novel coronavirus (SARS-CoV-2)"

Supplementary Table 1. Information of the 61 identified B-cell epitopes

| Protein | Start | End | Peptide | Length | Emini | Kolaskar | Selected for downstream analysis |
|---|---|---|---|---|---|---|---|
| Envelop | 5 | 11 | VSEETGT | 7 | 2.26 | 0.97 | |
| Envelop | 63 | 70 | KNLNSSRV | 8 | 3.471 | 1.002 | Yes |
| Membrane | 1 | 10 | MADSNGTITV | 10 | 0.56 | 0.977 | |
| Membrane | 109 | 115 | MWSFNPE | 7 | 1.405 | 0.93 | |
| Membrane | 161 | 171 | IKDLPKEITVA | 11 | 0.934 | 1.05 | |
| Membrane | 173 | 176 | SRTL | 4 | 1.504 | 1.011 | Yes |
| Membrane | 181 | 196 | LGASQRVAGDSGFAAY | 16 | 0.361 | 1.034 | |
| Membrane | 204 | 215 | YKLNTDHSSSSD | 12 | 7.078 | 0.993 | |
| Spike | 19 | 43 | TTRTQLPPAYTNSFTRGVYYPDKVF | 25 | 6.424 | 1.028 | Yes |
| Spike | 70 | 87 | VSGTNGTKRFDNPVLPFN | 18 | 1.184 | 0.994 | |
| Spike | 90 | 99 | VYFASTEKSN | 10 | 1.573 | 1.019 | Yes |
| Spike | 109 | 113 | TLDSK | 5 | 1.724 | 0.993 | |
| Spike | 146 | 154 | HKNNKSWME | 9 | 4.547 | 0.9 | |
| Spike | 158 | 166 | RVYSSANNC | 9 | 0.766 | 1.052 | |
| Spike | 180 | 187 | EGKQGNFK | 8 | 2.738 | 0.918 | |
| Spike | 206 | 209 | KHTP | 4 | 2.463 | 1.002 | Yes |
| Spike | 217 | 222 | PQGFSA | 6 | 0.809 | 1.02 | |
| Spike | 248 | 264 | YLTPGDSSSGWTAGAAA | 17 | 0.43 | 0.998 | |
| Spike | 280 | 288 | NENGTITDA | 9 | 1.46 | 0.909 | |
| Spike | 293 | 301 | LDPLSETKC | 9 | 0.844 | 1.06 | |
| Spike | 311 | 315 | GIYQT | 5 | 0.879 | 1.022 | |
| Spike | 317 | 327 | NFRVQPTESIV | 11 | 0.817 | 1.046 | |

| | | | | | | | |
|---|---|---|---|---|---|---|---|
| Spike | 380 | 387 | YGVSPTKL | 8 | 0.95 | 1.073 | |
| Spike | 405 | 430 | DEVRQIAPGQTGKIADYNYKLPDDFT | 26 | 5.81 | 1.001 | Yes |
| Spike | 438 | 448 | SNNLDSKVGGN | 11 | 1.294 | 0.957 | |
| Spike | 461 | 485 | LKPFERDISTEIYQAGSTPCNGVEG | 25 | 0.803 | 1.011 | |
| Spike | 494 | 507 | SYGFQPTNGVGYQP | 14 | 1.553 | 1.02 | Yes |
| Spike | 522 | 534 | ATVCGPKKSTNLV | 13 | 0.38 | 1.069 | |
| Spike | 545 | 559 | GLTGTGVLTESNKKF | 15 | 0.7 | 0.988 | |
| Spike | 564 | 582 | QFGRDIADTTDAVRDPQTL | 19 | 2.476 | 0.995 | |
| Spike | 590 | 593 | CSFG | 4 | 0.25 | 1.097 | |
| Spike | 595 | 607 | VSVITPGTNTSNQ | 13 | 0.803 | 1.013 | |
| Spike | 615 | 620 | VNCTEV | 6 | 0.309 | 1.119 | |
| Spike | 622 | 644 | VAIHADQLTPTWRVYSTGSNVFQ | 23 | 0.414 | 1.051 | |
| Spike | 655 | 667 | HVNNSYECDIPIG | 13 | 0.359 | 1.045 | |
| Spike | 671 | 688 | CASYQTQTNSPRRARSVA | 18 | 3.531 | 1.027 | Yes |
| Spike | 699 | 715 | LGAENSVAYSNNSIAIP | 17 | 0.312 | 1.026 | |
| Spike | 731 | 736 | MTKTSV | 6 | 1.067 | 0.995 | |
| Spike | 745 | 749 | DSTEC | 5 | 0.97 | 1.01 | |
| Spike | 771 | 782 | AVEQDKNTQEVF | 12 | 2.342 | 1.011 | Yes |
| Spike | 787 | 799 | QIYKTPPIKDFGG | 13 | 1.465 | 1.006 | Yes |
| Spike | 805 | 816 | ILPDPSKPSKRS | 12 | 4.69 | 1.019 | Yes |
| Spike | 833 | 836 | FIKQ | 4 | 0.853 | 1.047 | |
| Spike | 839 | 844 | DCLGDI | 6 | 0.223 | 1.07 | |
| Spike | 882 | 891 | ITSGWTFGAG | 10 | 0.187 | 0.965 | |
| Spike | 928 | 946 | NSAIGKIQDSLSSTASALG | 19 | 0.311 | 1.016 | |
| Spike | 950 | 958 | DVVNQNAQA | 9 | 0.974 | 1.038 | |
| Spike | 968 | 971 | SNFG | 4 | 0.749 | 0.938 | |

| | | | | | | | |
|---|---|---|---|---|---|---|---|
| Spike | 986 | 991 | KVEAEV | 6 | 0.869 | 1.077 | |
| Spike | 1018 | 1021 | IRAS | 4 | 0.754 | 1.025 | |
| Spike | 1023 | 1027 | NLAAT | 5 | 0.632 | 1.013 | |
| Spike | 1039 | 1042 | RVDF | 4 | 0.853 | 1.053 | |
| Spike | 1052 | 1058 | FPQSAPH | 7 | 1.381 | 1.059 | Yes |
| Spike | 1068 | 1091 | VPAQEKNFTTAPAICHDGKAHFPR | 24 | 1.063 | 1.03 | Yes |
| Spike | 1094 | 1097 | VFVS | 4 | 0.259 | 1.217 | |
| Spike | 1108 | 1123 | NFYEPQIITTDNTFVS | 16 | 1.039 | 1.007 | Yes |
| Spike | 1135 | 1151 | NTVYDPLQPELDSFKEE | 17 | 6.183 | 1.011 | Yes |
| Spike | 1153 | 1172 | DKYFKNHTSPDVDLGDISGI | 20 | 1.399 | 1.007 | Yes |
| Spike | 1190 | 1193 | AKNL | 4 | 1.087 | 1.005 | Yes |
| Spike | 1203 | 1209 | LGKYEQY | 7 | 2.512 | 1.035 | Yes |
| Spike | 1255 | 1265 | KFDEDDSEPVL | 11 | 2.654 | 1.003 | Yes |

Supplementary Table 2. Information of all identified T-cell epitopes

| Protein | Start | Length | Epi_AA | HLA_Count | HLA_Score | HLA_Types | Comments |
|---|---|---|---|---|---|---|---|
| Envelop | 4 | 9 | FVSEETGTL | 13 | 0.771 | HLA-A*02:06HLA-B*35:03,HLA-B*15:02,HLA-B*46:01,HLA-B*35:01,HLA-B*39:01HLA-C*07:02,HLA-C*15:02,HLA-C*03:04,HLA-C*04:01,HLA-C*03:03,HLA-C*06:02,HLA-C*12:03 | |
| Envelop | 5 | 9 | VSEETGTLI | 1 | 0.034 | HLA-C*15:02 | |
| Envelop | 6 | 9 | SEETGTLIV | 2 | 0.119 | HLA-B*40:01,HLA-B*40:02 | |
| Envelop | 10 | 9 | GTLIVNSVL | 1 | 0.034 | HLA-C*15:02 | |
| Envelop | 11 | 9 | TLIVNSVLL | 2 | 0.208 | HLA-A*02:07,HLA-A*02:01 | |
| Envelop | 12 | 9 | LIVNSVLLF | 6 | 0.399 | HLA-A*24:02,HLA-A*26:01HLA-B*46:01,HLA-B*15:02,HLA-B*15:01,HLA-B*35:01 | |
| Envelop | 13 | 9 | IVNSVLLFL | 7 | 0.484 | HLA-A*02:07,HLA-A*02:06,HLA-A*02:01HLA-C*15:02,HLA-C*03:03,HLA-C*03:04,HLA-C*12:03 | |
| Envelop | 15 | 9 | NSVLLFLAF | 3 | 0.166 | HLA-B*46:01,HLA-B*15:02,HLA-B*35:01 | |
| Envelop | 16 | 9 | SVLLFLAFV | 4 | 0.295 | HLA-A*02:03,HLA-A*02:07,HLA-A*02:06,HLA-A*02:01 | |
| Envelop | 17 | 9 | VLLFLAFVV | 3 | 0.260 | HLA-A*02:07,HLA-A*02:06,HLA-A*02:01 | |
| Envelop | 18 | 9 | LLFLAFVVF | 3 | 0.097 | HLA-A*32:01HLA-B*15:01,HLA-B*15:02 | |
| Envelop | 20 | 9 | FLAFVVFLL | 4 | 0.295 | HLA-A*02:03,HLA-A*02:07,HLA-A*02:06,HLA-A*02:01 | |
| Envelop | 21 | 9 | LAFVVFLLV | 3 | 0.106 | HLA-B*51:01,HLA-B*54:01HLA-C*12:03 | |
| Envelop | 23 | 9 | FVVFLLVTL | 7 | 0.440 | HLA-A*02:07,HLA-A*02:06HLA-B*46:01,HLA-B*35:03HLA-C*03:04,HLA-C*03:03,HLA-C*12:03 | |
| Envelop | 25 | 9 | VFLLVTLAI | 2 | 0.198 | HLA-A*24:02HLA-C*14:02 | |
| Envelop | 26 | 9 | FLLVTLAIL | 5 | 0.366 | HLA-A*02:03,HLA-A*02:07,HLA-A*02:06,HLA-A*02:01HLA-C*03:03 | |

| | | | | | | |
|---|---|---|---|---|---|---|
| Envelop | 29 | 9 | VTLAILTAL | 9 | 0.460 | HLA-A*02:06,HLA-A*32:01HLA-B*46:01,HLA-B*58:01,HLA-B*57:01HLA-C*15:02,HLA-C*03:03,HLA-C*03:04,HLA-C*12:03 |
| Envelop | 31 | 9 | LAILTALRL | 8 | 0.452 | HLA-B*46:01,HLA-B*51:01,HLA-B*58:01,HLA-B*35:03HLA-C*15:02,HLA-C*03:03,HLA-C*03:04,HLA-C*12:03 |
| Envelop | 34 | 9 | LTALRLCAY | 6 | 0.269 | HLA-A*01:01HLA-B*46:01,HLA-B*15:02,HLA-B*15:01,HLA-B*35:01HLA-C*12:03 |
| Envelop | 38 | 9 | RLCAYCCNI | 5 | 0.308 | HLA-A*32:01,HLA-A*02:03,HLA-A*02:07,HLA-A*02:06,HLA-A*02:01 |
| Envelop | 41 | 9 | AYCCNIVNV | 1 | 0.156 | HLA-A*24:02 |
| Envelop | 45 | 9 | NIVNVSLVK | 1 | 0.211 | HLA-A*11:01 |
| Envelop | 48 | 9 | NVSLVKPSF | 2 | 0.064 | HLA-B*35:01,HLA-B*15:02 |
| Envelop | 49 | 9 | VSLVKPSFY | 3 | 0.089 | HLA-B*58:01,HLA-B*57:01HLA-C*12:03 |
| Envelop | 50 | 9 | SLVKPSFYV | 4 | 0.295 | HLA-A*02:03,HLA-A*02:07,HLA-A*02:06,HLA-A*02:01 |
| Envelop | 51 | 9 | LVKPSFYVY | 6 | 0.247 | HLA-A*32:01HLA-B*46:01,HLA-B*15:02,HLA-B*15:01,HLA-B*35:01HLA-C*12:03 |
| Envelop | 55 | 9 | SFYVYSRVK | 1 | 0.054 | HLA-A*30:01 |
| Envelop | 57 | 9 | YVYSRVKNL | 10 | 0.614 | HLA-A*02:03,HLA-A*32:01HLA-C*07:02,HLA-C*15:02,HLA-C*03:04,HLA-C*04:01,HLA-C*14:02,HLA-C*03:03,HLA-C*06:02,HLA-C*12:03 |
| Envelop | 59 | 9 | YSRVKNLNS | 1 | 0.054 | HLA-A*30:01 |
| Envelop | 61 | 9 | RVKNLNSSR | 2 | 0.084 | HLA-A*30:01,HLA-A*03:01 |
| Envelop | 67 | 9 | SSRVPDLLV | 2 | 0.088 | HLA-A*30:01HLA-C*15:02 |
| Membrane | 6 | 9 | GTITVEELK | 1 | 0.211 | HLA-A*11:01 |
| Membrane | 7 | 9 | TITVEELKK | 1 | 0.211 | HLA-A*11:01 |

| | | | | | | |
|---|---|---|---|---|---|---|
| Membrane | 15 | 9 | KLLEQWNLV | 5 | 0.308 | HLA-A*32:01,HLA-A*02:03,HLA-A*02:07,HLA-A*02:06,HLA-A*02:01 |
| Membrane | 18 | 9 | EQWNLVIGF | 2 | 0.048 | HLA-A*32:01HLA-B*15:02 |
| Membrane | 21 | 9 | NLVIGFLFL | 2 | 0.208 | HLA-A*02:07,HLA-A*02:01 |
| Membrane | 22 | 9 | LVIGFLFLT | 2 | 0.175 | HLA-A*02:06,HLA-A*02:01 |
| Membrane | 23 | 9 | VIGFLFLTW | 4 | 0.238 | HLA-A*24:02,HLA-A*32:01HLA-B*58:01,HLA-B*57:01 |
| Membrane | 26 | 9 | FLFLTWICL | 2 | 0.208 | HLA-A*02:07,HLA-A*02:01 |
| Membrane | 27 | 9 | LFLTWICLL | 1 | 0.042 | HLA-C*14:02 |
| Membrane | 29 | 9 | LTWICLLQF | 3 | 0.082 | HLA-A*32:01HLA-B*58:01,HLA-B*57:01 |
| Membrane | 31 | 9 | WICLLQFAY | 1 | 0.029 | HLA-B*35:01 |
| Membrane | 37 | 9 | FAYANRNRF | 16 | 0.898 | HLA-A*32:01HLA-B*15:01,HLA-B*35:03,HLA-B*58:01,HLA-B*15:02,HLA-B*46:01,HLA-B*35:01,HLA-B*51:01,HLA-B*57:01HLA-C*07:02,HLA-C*03:04,HLA-C*04:01,HLA-C*14:02,HLA-C*03:03,HLA-C*06:02,HLA-C*12:03 |
| Membrane | 38 | 9 | AYANRNRFL | 5 | 0.499 | HLA-A*24:02HLA-C*07:02,HLA-C*14:02,HLA-C*06:02,HLA-C*04:01 |
| Membrane | 39 | 9 | YANRNRFLY | 12 | 0.663 | HLA-A*01:01,HLA-A*32:01HLA-B*15:01,HLA-B*58:01,HLA-B*15:02,HLA-B*46:01,HLA-B*35:01,HLA-B*57:01HLA-C*07:02,HLA-C*03:03,HLA-C*06:02,HLA-C*12:03 |
| Membrane | 40 | 9 | ANRNRFLYI | 1 | 0.054 | HLA-A*30:01 |
| Membrane | 41 | 9 | NRNRFLYII | 3 | 0.259 | HLA-B*39:01HLA-C*07:02,HLA-C*06:02 |
| Membrane | 42 | 9 | RNRFLYIIK | 1 | 0.054 | HLA-A*30:01 |
| Membrane | 43 | 9 | NRFLYIIKL | 4 | 0.301 | HLA-B*39:01HLA-C*07:02,HLA-C*14:02,HLA-C*06:02 |
| Membrane | 44 | 9 | RFLYIIKLI | 2 | 0.168 | HLA-A*24:02,HLA-A*32:01 |

| | | | | | | |
|---|---|---|---|---|---|---|
| Membrane | 45 | 9 | FLYIIKLIF | 2 | 0.062 | HLA-A*32:01HLA-B*15:01 |
| Membrane | 46 | 9 | LYIIKLIFL | 3 | 0.350 | HLA-A*24:02HLA-C*07:02,HLA-C*14:02 |
| Membrane | 47 | 9 | YIIKLIFLW | 4 | 0.238 | HLA-A*24:02,HLA-A*32:01HLA-B*58:01,HLA-B*57:01 |
| Membrane | 50 | 9 | KLIFLWLLW | 4 | 0.238 | HLA-A*24:02,HLA-A*32:01HLA-B*58:01,HLA-B*57:01 |
| Membrane | 53 | 9 | FLWLLWPVT | 3 | 0.260 | HLA-A*02:07,HLA-A*02:06,HLA-A*02:01 |
| Membrane | 54 | 9 | LWLLWPVTL | 3 | 0.350 | HLA-A*24:02HLA-C*07:02,HLA-C*14:02 |
| Membrane | 55 | 9 | WLLWPVTLA | 4 | 0.295 | HLA-A*02:03,HLA-A*02:07,HLA-A*02:06,HLA-A*02:01 |
| Membrane | 56 | 9 | LLWPVTLAC | 1 | 0.085 | HLA-A*02:07 |
| Membrane | 57 | 9 | LWPVTLACF | 2 | 0.198 | HLA-A*24:02HLA-C*14:02 |
| Membrane | 58 | 9 | WPVTLACFV | 4 | 0.126 | HLA-B*35:01,HLA-B*51:01,HLA-B*54:01,HLA-B*35:03 |
| Membrane | 60 | 9 | VTLACFVLA | 1 | 0.052 | HLA-A*02:06 |
| Membrane | 61 | 9 | TLACFVLAA | 4 | 0.295 | HLA-A*02:03,HLA-A*02:07,HLA-A*02:06,HLA-A*02:01 |
| Membrane | 62 | 9 | LACFVLAAV | 6 | 0.329 | HLA-A*02:06HLA-B*51:01,HLA-B*54:01HLA-C*03:04,HLA-C*03:03,HLA-C*12:03 |
| Membrane | 65 | 9 | FVLAAVYRI | 8 | 0.497 | HLA-A*32:01,HLA-A*02:03,HLA-A*02:07,HLA-A*02:06,HLA-A*02:01HLA-B*51:01HLA-C*15:02,HLA-C*03:04 |
| Membrane | 67 | 9 | LAAVYRINW | 3 | 0.082 | HLA-A*32:01HLA-B*58:01,HLA-B*57:01 |
| Membrane | 68 | 9 | AAVYRINWI | 4 | 0.214 | HLA-C*15:02,HLA-C*03:03,HLA-C*06:02,HLA-C*12:03 |
| Membrane | 71 | 9 | YRINWITGG | 1 | 0.152 | HLA-C*07:02 |
| Membrane | 72 | 9 | RINWITGGI | 2 | 0.047 | HLA-A*32:01HLA-C*15:02 |
| Membrane | 76 | 9 | ITGGIAIAM | 7 | 0.390 | HLA-B*46:01,HLA-B*15:02,HLA-B*35:01HLA-C*15:02,HLA-C*03:03,HLA-C*03:04,HLA-C*12:03 |
| Membrane | 80 | 9 | IAIAMACLV | 7 | 0.322 | HLA-A*02:06HLA-B*58:01,HLA-B*51:01,HLA-B*54:01HLA-C*15:02,HLA-C*03:03,HLA-C*12:03 |

| | | | | | | |
|---|---|---|---|---|---|---|
| Membrane | 82 | 9 | IAMACLVGL | 10 | 0.599 | HLA-A*02:06,HLA-A*02:01HLA-B*46:01,HLA-B*58:01,HLA-B*35:03,HLA-B*35:01HLA-C*15:02,HLA-C*03:03,HLA-C*03:04,HLA-C*12:03 |
| Membrane | 83 | 9 | AMACLVGLM | 6 | 0.430 | HLA-A*02:03,HLA-A*02:07,HLA-A*02:01HLA-B*46:01,HLA-B*15:01,HLA-B*15:02 |
| Membrane | 84 | 9 | MACLVGLMW | 3 | 0.082 | HLA-A*32:01HLA-B*58:01,HLA-B*57:01 |
| Membrane | 87 | 9 | LVGLMWLSY | 3 | 0.098 | HLA-A*01:01HLA-B*35:01,HLA-B*15:02 |
| Membrane | 89 | 9 | GLMWLSYFI | 5 | 0.308 | HLA-A*32:01,HLA-A*02:03,HLA-A*02:07,HLA-A*02:06,HLA-A*02:01 |
| Membrane | 90 | 9 | LMWLSYFIA | 3 | 0.260 | HLA-A*02:07,HLA-A*02:06,HLA-A*02:01 |
| Membrane | 92 | 9 | WLSYFIASF | 4 | 0.200 | HLA-A*32:01HLA-B*46:01,HLA-B*15:01,HLA-B*15:02 |
| Membrane | 93 | 9 | LSYFIASFR | 1 | 0.211 | HLA-A*11:01 |
| Membrane | 94 | 9 | SYFIASFRL | 4 | 0.439 | HLA-A*24:02HLA-C*07:02,HLA-C*14:02,HLA-C*06:02 |
| Membrane | 95 | 9 | YFIASFRLF | 8 | 0.625 | HLA-A*24:02HLA-B*46:01,HLA-B*15:02,HLA-B*15:01,HLA-B*35:01HLA-C*07:02,HLA-C*14:02,HLA-C*04:01 |
| Membrane | 96 | 9 | FIASFRLFA | 4 | 0.240 | HLA-A*02:03,HLA-A*02:06,HLA-A*02:01HLA-B*54:01 |
| Membrane | 99 | 9 | SFRLFARTR | 1 | 0.054 | HLA-A*30:01 |
| Membrane | 101 | 9 | RLFARTRSM | 13 | 0.741 | HLA-A*02:03,HLA-A*30:01,HLA-A*32:01HLA-B*15:01,HLA-B*15:02,HLA-B*07:02HLA-C*07:02,HLA-C*03:04,HLA-C*04:01,HLA-C*14:02,HLA-C*03:03,HLA-C*06:02,HLA-C*12:03 |
| Membrane | 102 | 9 | LFARTRSMW | 2 | 0.198 | HLA-A*24:02HLA-C*14:02 |
| Membrane | 104 | 9 | ARTRSMWSF | 2 | 0.241 | HLA-C*07:02,HLA-C*06:02 |
| Membrane | 105 | 9 | RTRSMWSFN | 1 | 0.054 | HLA-A*30:01 |
| Membrane | 107 | 9 | RSMWSFNPE | 1 | 0.054 | HLA-A*30:01 |

| | | | | | | |
|---|---|---|---|---|---|---|
| Membrane | 108 | 9 | SMWSFNPET | 1 | 0.123 | HLA-A*02:01 |
| Membrane | 110 | 9 | WSFNPETNI | 5 | 0.268 | HLA-B*58:01,HLA-B*51:01HLA-C*15:02,HLA-C*03:04,HLA-C*12:03 |
| Membrane | 111 | 9 | SFNPETNIL | 5 | 0.499 | HLA-A*24:02HLA-C*07:02,HLA-C*14:02,HLA-C*06:02,HLA-C*04:01 |
| Membrane | 112 | 9 | FNPETNILL | 2 | 0.211 | HLA-C*07:02,HLA-C*04:01 |
| Membrane | 116 | 9 | TNILLNVPL | 1 | 0.018 | HLA-B*39:01 |
| Membrane | 122 | 9 | VPLHGTILT | 1 | 0.030 | HLA-B*54:01 |
| Membrane | 125 | 9 | HGTILTRPL | 1 | 0.100 | HLA-C*03:04 |
| Membrane | 126 | 9 | GTILTRPLL | 1 | 0.034 | HLA-C*15:02 |
| Membrane | 130 | 9 | TRPLLESEL | 3 | 0.259 | HLA-B*39:01HLA-C*07:02,HLA-C*06:02 |
| Membrane | 131 | 9 | RPLLESELV | 2 | 0.078 | HLA-B*51:01,HLA-B*07:02 |
| Membrane | 134 | 9 | LESELVIGA | 1 | 0.019 | HLA-B*40:02 |
| Membrane | 136 | 9 | SELVIGAVI | 3 | 0.145 | HLA-B*44:03,HLA-B*40:01,HLA-B*40:02 |
| Membrane | 138 | 9 | LVIGAVILR | 1 | 0.211 | HLA-A*11:01 |
| Membrane | 142 | 9 | AVILRGHLR | 1 | 0.211 | HLA-A*11:01 |
| Membrane | 144 | 9 | ILRGHLRIA | 2 | 0.089 | HLA-A*02:03,HLA-A*30:01 |
| Membrane | 148 | 9 | HLRIAGHHL | 4 | 0.161 | HLA-A*30:01HLA-B*15:01,HLA-B*15:02,HLA-B*07:02 |
| Membrane | 150 | 9 | RIAGHHLGR | 2 | 0.241 | HLA-A*11:01,HLA-A*03:01 |
| Membrane | 164 | 9 | LPKEITVAT | 4 | 0.092 | HLA-B*35:01,HLA-B*07:02,HLA-B*54:01,HLA-B*35:03 |
| Membrane | 168 | 9 | ITVATSRTL | 6 | 0.337 | HLA-B*46:01,HLA-B*57:01HLA-C*15:02,HLA-C*03:03,HLA-C*03:04,HLA-C*12:03 |
| Membrane | 170 | 9 | VATSRTLSY | 10 | 0.639 | HLA-A*01:01HLA-B*15:01,HLA-B*58:01,HLA-B*15:02,HLA-B*46:01,HLA-B*35:01HLA-C*07:02,HLA-C*03:03,HLA-C*06:02,HLA-C*12:03 |

| | | | | | | |
|---|---|---|---|---|---|---|
| Membrane | 171 | 9 | ATSRTLSYY | 8 | 0.473 | HLA-A*11:01,HLA-A*01:01,HLA-A*30:01HLA-B*58:01,HLA-B*15:01,HLA-B*15:02,HLA-B*57:01HLA-C*12:03 |
| Membrane | 172 | 9 | TSRTLSYYK | 3 | 0.296 | HLA-A*11:01,HLA-A*30:01,HLA-A*03:01 |
| Membrane | 173 | 9 | SRTLSYYKL | 3 | 0.259 | HLA-B*39:01HLA-C*07:02,HLA-C*06:02 |
| Membrane | 174 | 9 | RTLSYYKLG | 1 | 0.054 | HLA-A*30:01 |
| Membrane | 179 | 9 | YKLGASQRV | 1 | 0.089 | HLA-C*06:02 |
| Membrane | 188 | 9 | AGDSGFAAY | 3 | 0.122 | HLA-A*01:01HLA-B*35:01HLA-C*04:01 |
| Membrane | 191 | 9 | SGFAAYSRY | 7 | 0.475 | HLA-B*46:01,HLA-B*15:02,HLA-B*15:01,HLA-B*35:01HLA-C*07:02,HLA-C*06:02,HLA-C*12:03 |
| Membrane | 193 | 9 | FAAYSRYRI | 6 | 0.369 | HLA-B*51:01HLA-C*15:02,HLA-C*03:04,HLA-C*03:03,HLA-C*06:02,HLA-C*12:03 |
| Membrane | 196 | 9 | YSRYRIGNY | 6 | 0.447 | HLA-B*46:01,HLA-B*15:01,HLA-B*15:02HLA-C*07:02,HLA-C*06:02,HLA-C*12:03 |
| Membrane | 198 | 9 | RYRIGNYKL | 5 | 0.464 | HLA-A*24:02,HLA-A*30:01HLA-C*07:02,HLA-C*14:02,HLA-C*04:01 |
| Membrane | 209 | 9 | DHSSSSDNI | 1 | 0.018 | HLA-B*39:01 |
| Membrane | 211 | 9 | SSSSDNIAL | 3 | 0.205 | HLA-C*15:02,HLA-C*03:03,HLA-C*03:04 |
| Membrane | 212 | 9 | SSSDNIALL | 6 | 0.465 | HLA-C*07:02,HLA-C*15:02,HLA-C*03:04,HLA-C*03:03,HLA-C*06:02,HLA-C*12:03 |
| Membrane | 213 | 9 | SSDNIALLV | 3 | 0.087 | HLA-A*01:01HLA-C*15:02,HLA-C*12:03 |
| Spike | 2 | 9 | FVFLVLLPL | 7 | 0.552 | HLA-A*02:07,HLA-A*02:06,HLA-A*02:01HLA-B*46:01HLA-C*03:04,HLA-C*03:03,HLA-C*12:03 |
| Spike | 8 | 9 | LPLVSSQCV | 5 | 0.148 | HLA-B*35:03,HLA-B*54:01,HLA-B*07:02,HLA-B*35:01,HLA-B*51:01 |
| Spike | 10 | 9 | LVSSQCVNL | 2 | 0.171 | HLA-C*03:04,HLA-C*03:03 |

| | | | | | | |
|---|---|---|---|---|---|---|
| Spike | 19 | 9 | TTRTQLPPA | 1 | 0.054 | HLA-A*30:01 |
| Spike | 20 | 9 | TRTQLPPAY | 2 | 0.241 | HLA-C*07:02,HLA-C*06:02 |
| Spike | 21 | 9 | RTQLPPAYT | 1 | 0.054 | HLA-A*30:01 |
| Spike | 24 | 9 | LPPAYTNSF | 5 | 0.178 | HLA-B*35:01,HLA-B*51:01,HLA-B*35:03,HLA-B*07:02HLA-C*04:01 |
| Spike | 28 | 9 | YTNSFTRGV | 6 | 0.329 | HLA-A*02:03,HLA-A*02:06HLA-C*15:02,HLA-C*06:02,HLA-C*03:04,HLA-C*12:03 |
| Spike | 29 | 9 | TNSFTRGVY | 1 | 0.035 | HLA-B*15:02 |
| Spike | 30 | 9 | NSFTRGVYY | 7 | 0.569 | HLA-A*11:01,HLA-A*01:01HLA-B*35:01,HLA-B*15:02HLA-C*07:02,HLA-C*06:02,HLA-C*12:03 |
| Spike | 35 | 9 | GVYYPDKVF | 3 | 0.097 | HLA-A*32:01HLA-B*15:01,HLA-B*15:02 |
| Spike | 37 | 9 | YYPDKVFRS | 1 | 0.152 | HLA-C*07:02 |
| Spike | 38 | 9 | YPDKVFRSS | 2 | 0.052 | HLA-B*54:01,HLA-B*07:02 |
| Spike | 41 | 9 | KVFRSSVLH | 3 | 0.296 | HLA-A*11:01,HLA-A*30:01,HLA-A*03:01 |
| Spike | 42 | 9 | VFRSSVLHS | 1 | 0.054 | HLA-A*30:01 |
| Spike | 43 | 9 | FRSSVLHST | 2 | 0.169 | HLA-B*39:01HLA-C*07:02 |
| Spike | 46 | 9 | SVLHSTQDL | 2 | 0.171 | HLA-C*03:04,HLA-C*03:03 |
| Spike | 47 | 9 | VLHSTQDLF | 4 | 0.253 | HLA-A*24:02,HLA-A*32:01HLA-B*15:01,HLA-B*15:02 |
| Spike | 48 | 9 | LHSTQDLFL | 2 | 0.169 | HLA-B*39:01HLA-C*07:02 |
| Spike | 50 | 9 | STQDLFLPF | 10 | 0.651 | HLA-A*24:02,HLA-A*26:01,HLA-A*32:01HLA-B*35:01,HLA-B*15:01,HLA-B*15:02HLA-C*07:02,HLA-C*03:04,HLA-C*03:03,HLA-C*12:03 |
| Spike | 51 | 9 | TQDLFLPFF | 1 | 0.060 | HLA-C*04:01 |
| Spike | 55 | 9 | FLPFFSNVT | 2 | 0.120 | HLA-A*02:03,HLA-A*02:07 |
| Spike | 56 | 9 | LPFFSNVTW | 7 | 0.218 | HLA-B*35:03,HLA-B*58:01,HLA-B*54:01,HLA-B*07:02,HLA-B*35:01,HLA-B*51:01,HLA-B*57:01 |

| | | | | | | |
|---|---|---|---|---|---|---|
| Spike | 57 | 9 | PFFSNVTWF | 1 | 0.156 | HLA-A*24:02 |
| Spike | 59 | 9 | FSNVTWFHA | 1 | 0.030 | HLA-B*54:01 |
| Spike | 60 | 9 | SNVTWFHAI | 2 | 0.047 | HLA-A*32:01HLA-C*15:02 |
| Spike | 62 | 9 | VTWFHAIHV | 4 | 0.228 | HLA-A*02:06,HLA-A*02:01HLA-C*15:02,HLA-C*12:03 |
| Spike | 65 | 9 | FHAIHVSGT | 1 | 0.018 | HLA-B*39:01 |
| Spike | 69 | 9 | HVSGTNGTK | 2 | 0.241 | HLA-A*11:01,HLA-A*03:01 |
| Spike | 75 | 9 | GTKRFDNPV | 1 | 0.054 | HLA-A*30:01 |
| Spike | 78 | 9 | RFDNPVLPF | 5 | 0.422 | HLA-A*24:02,HLA-A*32:01HLA-C*07:02,HLA-C*14:02,HLA-C*04:01 |
| Spike | 81 | 9 | NPVLPFNDG | 1 | 0.030 | HLA-B*54:01 |
| Spike | 83 | 9 | VLPFNDGVY | 3 | 0.187 | HLA-B*46:01,HLA-B*15:01,HLA-B*15:02 |
| Spike | 84 | 9 | LPFNDGVYF | 8 | 0.274 | HLA-B*35:03,HLA-B*15:02,HLA-B*54:01,HLA-B*07:02,HLA-B*35:01,HLA-B*51:01HLA-C*03:03,HLA-C*12:03 |
| Spike | 89 | 9 | GVYFASTEK | 3 | 0.296 | HLA-A*11:01,HLA-A*30:01,HLA-A*03:01 |
| Spike | 92 | 9 | FASTEKSNI | 5 | 0.280 | HLA-B*51:01HLA-C*15:02,HLA-C*03:03,HLA-C*03:04,HLA-C*12:03 |
| Spike | 93 | 9 | ASTEKSNII | 1 | 0.034 | HLA-C*15:02 |
| Spike | 97 | 9 | KSNIIRGWI | 4 | 0.158 | HLA-A*30:01HLA-B*58:01,HLA-B*57:01HLA-C*15:02 |
| Spike | 102 | 9 | RGWIFGTTL | 3 | 0.184 | HLA-A*32:01HLA-C*03:04,HLA-C*03:03 |
| Spike | 109 | 9 | TLDSKTQSL | 4 | 0.314 | HLA-A*02:07HLA-B*39:01HLA-C*07:02,HLA-C*04:01 |
| Spike | 113 | 9 | KTQSLLIVN | 1 | 0.054 | HLA-A*30:01 |
| Spike | 118 | 9 | LIVNNATNV | 3 | 0.206 | HLA-A*02:03HLA-C*03:04,HLA-C*03:03 |
| Spike | 119 | 9 | IVNNATNVV | 3 | 0.153 | HLA-C*15:02,HLA-C*03:04,HLA-C*12:03 |
| Spike | 125 | 9 | NVVIKVCEF | 2 | 0.063 | HLA-A*26:01HLA-B*15:02 |
| Spike | 127 | 9 | VIKVCEFQF | 1 | 0.013 | HLA-A*32:01 |

| | | | | | | |
|---|---|---|---|---|---|---|
| Spike | 132 | 9 | EFQFCNDPF | 2 | 0.198 | HLA-A*24:02HLA-C*14:02 |
| Spike | 133 | 9 | FQFCNDPFL | 11 | 0.808 | HLA-A*02:06,HLA-A*02:01HLA-B*40:01,HLA-B*15:01,HLA-B*15:02,HLA-B*39:01HLA-C*07:02,HLA-C*03:04,HLA-C*03:03,HLA-C*06:02,HLA-C*12:03 |
| Spike | 135 | 9 | FCNDPFLGV | 2 | 0.086 | HLA-A*02:06HLA-C*15:02 |
| Spike | 142 | 9 | GVYYHKNNK | 3 | 0.296 | HLA-A*11:01,HLA-A*30:01,HLA-A*03:01 |
| Spike | 144 | 9 | YYHKNNKSW | 4 | 0.439 | HLA-A*24:02HLA-C*07:02,HLA-C*14:02,HLA-C*06:02 |
| Spike | 145 | 9 | YHKNNKSWM | 2 | 0.241 | HLA-C*07:02,HLA-C*06:02 |
| Spike | 150 | 9 | KSWMESEFR | 1 | 0.211 | HLA-A*11:01 |
| Spike | 151 | 9 | SWMESEFRV | 1 | 0.156 | HLA-A*24:02 |
| Spike | 152 | 9 | WMESEFRVY | 4 | 0.173 | HLA-B*35:01,HLA-B*15:01,HLA-B*15:02HLA-C*04:01 |
| Spike | 155 | 9 | SEFRVYSSA | 1 | 0.019 | HLA-B*40:02 |
| Spike | 158 | 9 | RVYSSANNC | 1 | 0.054 | HLA-A*30:01 |
| Spike | 160 | 9 | YSSANNCTF | 10 | 0.584 | HLA-B*15:01,HLA-B*58:01,HLA-B*15:02,HLA-B*35:01,HLA-B*57:01HLA-C*07:02,HLA-C*03:04,HLA-C*04:01,HLA-C*03:03,HLA-C*12:03 |
| Spike | 162 | 9 | SANNCTFEY | 9 | 0.690 | HLA-A*11:01,HLA-A*01:01HLA-B*15:01,HLA-B*58:01,HLA-B*15:02,HLA-B*46:01,HLA-B*35:01HLA-C*07:02,HLA-C*12:03 |
| Spike | 167 | 9 | TFEYVSQPF | 5 | 0.438 | HLA-A*24:02HLA-B*35:01HLA-C*07:02,HLA-C*14:02,HLA-C*04:01 |
| Spike | 168 | 9 | FEYVSQPFL | 3 | 0.179 | HLA-B*40:01,HLA-B*40:02HLA-C*04:01 |
| Spike | 169 | 9 | EYVSQPFLM | 4 | 0.439 | HLA-A*24:02HLA-C*07:02,HLA-C*14:02,HLA-C*06:02 |
| Spike | 171 | 9 | VSQPFLMDL | 6 | 0.386 | HLA-B*57:01HLA-C*07:02,HLA-C*03:04,HLA-C*15:02,HLA-C*03:03,HLA-C*12:03 |
| Spike | 186 | 9 | FKNLREFVF | 1 | 0.152 | HLA-C*07:02 |

| | | | | | | |
|---|---|---|---|---|---|---|
| Spike | 187 | 9 | KNLREFVFK | 2 | 0.266 | HLA-A*11:01,HLA-A*30:01 |
| Spike | 189 | 9 | LREFVFKNI | 1 | 0.089 | HLA-C*06:02 |
| Spike | 190 | 9 | REFVFKNID | 1 | 0.019 | HLA-B*40:02 |
| Spike | 192 | 9 | FVFKNIDGY | 10 | 0.627 | HLA-A*26:01HLA-B*46:01,HLA-B*15:02,HLA-B*15:01,HLA-B*35:01HLA-C*07:02,HLA-C*03:04,HLA-C*14:02,HLA-C*03:03,HLA-C*12:03 |
| Spike | 193 | 9 | VFKNIDGYF | 4 | 0.409 | HLA-A*24:02HLA-C*07:02,HLA-C*14:02,HLA-C*04:01 |
| Spike | 195 | 9 | KNIDGYFKI | 1 | 0.013 | HLA-A*32:01 |
| Spike | 196 | 9 | NIDGYFKIY | 3 | 0.098 | HLA-A*01:01HLA-B*35:01,HLA-B*15:02 |
| Spike | 202 | 9 | KIYSKHTPI | 6 | 0.408 | HLA-A*02:01,HLA-A*02:07,HLA-A*30:01,HLA-A*32:01HLA-C*15:02,HLA-C*03:04 |
| Spike | 204 | 9 | YSKHTPINL | 7 | 0.567 | HLA-B*46:01HLA-C*07:02,HLA-C*15:02,HLA-C*03:04,HLA-C*03:03,HLA-C*06:02,HLA-C*12:03 |
| Spike | 205 | 9 | SKHTPINLV | 1 | 0.089 | HLA-C*06:02 |
| Spike | 208 | 9 | TPINLVRDL | 3 | 0.089 | HLA-B*51:01,HLA-B*35:03,HLA-B*07:02 |
| Spike | 212 | 9 | LVRDLPQGF | 3 | 0.187 | HLA-B*46:01,HLA-B*15:01,HLA-B*15:02 |
| Spike | 215 | 9 | DLPQGFSAL | 1 | 0.028 | HLA-A*26:01 |
| Spike | 216 | 9 | LPQGFSALE | 1 | 0.030 | HLA-B*54:01 |
| Spike | 218 | 9 | QGFSALEPL | 1 | 0.100 | HLA-C*03:04 |
| Spike | 221 | 9 | SALEPLVDL | 5 | 0.235 | HLA-B*35:03HLA-C*15:02,HLA-C*03:03,HLA-C*03:04,HLA-C*12:03 |
| Spike | 223 | 9 | LEPLVDLPI | 2 | 0.119 | HLA-B*40:01,HLA-B*40:02 |
| Spike | 229 | 9 | LPIGINITR | 2 | 0.059 | HLA-B*35:01,HLA-B*54:01 |
| Spike | 235 | 9 | ITRFQTLLA | 1 | 0.054 | HLA-A*30:01 |
| Spike | 236 | 9 | TRFQTLLAL | 4 | 0.318 | HLA-B*39:01HLA-C*07:02,HLA-C*06:02,HLA-C*04:01 |
| Spike | 240 | 9 | TLLALHRSY | 4 | 0.126 | HLA-A*32:01HLA-B*35:01,HLA-B*15:01,HLA-B*15:02 |

| | | | | | | |
|---|---|---|---|---|---|---|
| Spike | 241 | 9 | LLALHRSYL | 7 | 0.525 | HLA-A*02:03,HLA-A*02:07,HLA-A*02:01HLA-B*07:02HLA-C*03:04,HLA-C*03:03,HLA-C*06:02 |
| Spike | 258 | 9 | WTAGAAAYY | 10 | 0.565 | HLA-A*01:01,HLA-A*26:01HLA-B*58:01,HLA-B*35:01,HLA-B*15:01,HLA-B*15:02HLA-C*07:02,HLA-C*03:03,HLA-C*06:02,HLA-C*12:03 |
| Spike | 259 | 9 | TAGAAAYYV | 4 | 0.161 | HLA-A*02:06HLA-B*51:01HLA-C*15:02,HLA-C*12:03 |
| Spike | 261 | 9 | GAAAYYVGY | 5 | 0.344 | HLA-A*11:01HLA-B*35:01,HLA-B*15:01,HLA-B*15:02HLA-C*12:03 |
| Spike | 262 | 9 | AAAYYVGYL | 7 | 0.567 | HLA-B*46:01HLA-C*07:02,HLA-C*15:02,HLA-C*03:04,HLA-C*03:03,HLA-C*06:02,HLA-C*12:03 |
| Spike | 265 | 9 | YYVGYLQPR | 1 | 0.152 | HLA-C*07:02 |
| Spike | 267 | 9 | VGYLQPRTF | 4 | 0.362 | HLA-B*46:01HLA-C*07:02,HLA-C*06:02,HLA-C*12:03 |
| Spike | 268 | 9 | GYLQPRTFL | 3 | 0.350 | HLA-A*24:02HLA-C*07:02,HLA-C*14:02 |
| Spike | 269 | 9 | YLQPRTFLL | 13 | 0.996 | HLA-A*32:01,HLA-A*24:02,HLA-A*02:03,HLA-A*02:07,HLA-A*02:06,HLA-A*02:01HLA-C*07:02,HLA-C*03:04,HLA-C*04:01,HLA-C*14:02,HLA-C*03:03,HLA-C*06:02,HLA-C*12:03 |
| Spike | 270 | 9 | LQPRTFLLK | 2 | 0.241 | HLA-A*11:01,HLA-A*03:01 |
| Spike | 271 | 9 | QPRTFLLKY | 1 | 0.029 | HLA-B*35:01 |
| Spike | 285 | 9 | ITDAVDCAL | 6 | 0.318 | HLA-A*01:01HLA-C*15:02,HLA-C*03:04,HLA-C*04:01,HLA-C*03:03,HLA-C*12:03 |
| Spike | 288 | 9 | AVDCALDPL | 4 | 0.283 | HLA-A*02:06HLA-C*03:04,HLA-C*03:03,HLA-C*04:01 |
| Spike | 292 | 9 | ALDPLSETK | 2 | 0.241 | HLA-A*11:01,HLA-A*03:01 |
| Spike | 296 | 9 | LSETKCTLK | 1 | 0.211 | HLA-A*11:01 |
| Spike | 298 | 9 | ETKCTLKSF | 2 | 0.063 | HLA-A*26:01HLA-B*15:02 |
| Spike | 302 | 9 | TLKSFTVEK | 3 | 0.296 | HLA-A*11:01,HLA-A*30:01,HLA-A*03:01 |

| | | | | | | | |
|---|---|---|---|---|---|---|---|
| Spike | 304 | 9 | KSFTVEKGI | 4 | 0.116 | HLA-A*32:01HLA-B*58:01,HLA-B*57:01HLA-C*15:02 | |
| Spike | 310 | 9 | KGIYQTSNF | 4 | 0.175 | HLA-A*32:01HLA-B*46:01,HLA-B*15:01,HLA-B*57:01 | |
| Spike | 311 | 9 | GIYQTSNFR | 2 | 0.241 | HLA-A*11:01,HLA-A*03:01 | |
| Spike | 312 | 9 | IYQTSNFRV | 3 | 0.350 | HLA-A*24:02HLA-C*07:02,HLA-C*14:02 | |
| Spike | 318 | 9 | FRVQPTESI | 3 | 0.259 | HLA-B*39:01HLA-C*07:02,HLA-C*06:02 | |
| Spike | 319 | 9 | RVQPTESIV | 1 | 0.034 | HLA-C*15:02 | |
| Spike | 321 | 9 | QPTESIVRF | 2 | 0.040 | HLA-B*35:01,HLA-B*35:03 | |
| Spike | 324 | 9 | ESIVRFPNI | 2 | 0.084 | HLA-A*26:01HLA-B*51:01 | |
| Spike | 326 | 9 | IVRFPNITN | 1 | 0.054 | HLA-A*30:01 | |
| Spike | 327 | 9 | VRFPNITNL | 5 | 0.320 | HLA-B*39:01HLA-C*07:02,HLA-C*14:02,HLA-C*06:02,HLA-C*12:03 | |
| Spike | 329 | 9 | FPNITNLCP | 1 | 0.030 | HLA-B*54:01 | |
| Spike | 334 | 9 | NLCPFGEVF | 2 | 0.085 | HLA-B*15:01,HLA-B*15:02 | |
| Spike | 336 | 9 | CPFGEVFNA | 2 | 0.059 | HLA-B*35:01,HLA-B*54:01 | |
| Spike | 339 | 9 | GEVFNATRF | 3 | 0.145 | HLA-B*44:03,HLA-B*40:01,HLA-B*40:02 | |
| Spike | 340 | 9 | EVFNATRFA | 2 | 0.058 | HLA-A*26:01HLA-B*54:01 | |
| Spike | 342 | 9 | FNATRFASV | 1 | 0.052 | HLA-A*02:06 | |
| Spike | 343 | 9 | NATRFASVY | 4 | 0.185 | HLA-B*46:01,HLA-B*15:02,HLA-B*35:01HLA-C*12:03 | |
| Spike | 344 | 9 | ATRFASVYA | 1 | 0.054 | HLA-A*30:01 | |
| Spike | 345 | 9 | TRFASVYAW | 4 | 0.271 | HLA-A*32:01HLA-B*39:01HLA-C*07:02,HLA-C*06:02 | |
| Spike | 348 | 9 | ASVYAWNRK | 2 | 0.266 | HLA-A*11:01,HLA-A*30:01 | RBD |
| Spike | 349 | 9 | SVYAWNRKR | 2 | 0.241 | HLA-A*11:01,HLA-A*03:01 | RBD |
| Spike | 350 | 9 | VYAWNRKRI | 3 | 0.350 | HLA-A*24:02HLA-C*07:02,HLA-C*14:02 | RBD |
| Spike | 354 | 9 | NRKRISNCV | 1 | 0.089 | HLA-C*06:02 | RBD |
| Spike | 355 | 9 | RKRISNCVA | 1 | 0.054 | HLA-A*30:01 | RBD |
| Spike | 357 | 9 | RISNCVADY | 4 | 0.127 | HLA-A*03:01,HLA-A*32:01HLA-B*15:01,HLA-B*15:02 | RBD |

| | | | | | | | |
|---|---|---|---|---|---|---|---|
| Spike | 361 | 9 | CVADYSVLY | 9 | 0.647 | HLA-A*11:01,HLA-A*01:01,HLA-A*26:01HLA-B*35:01,HLA-B*15:01,HLA-B*15:02HLA-C*07:02,HLA-C*06:02,HLA-C*12:03 | RBD |
| Spike | 366 | 9 | SVLYNSASF | 6 | 0.297 | HLA-A*32:01HLA-B*35:01,HLA-B*15:01,HLA-B*15:02HLA-C*03:04,HLA-C*03:03 | RBD |
| Spike | 369 | 9 | YNSASFSTF | 8 | 0.602 | HLA-A*24:02HLA-B*46:01,HLA-B*15:02,HLA-B*15:01,HLA-B*35:01HLA-C*03:04,HLA-C*03:03,HLA-C*04:01 | RBD |
| Spike | 370 | 9 | NSASFSTFK | 3 | 0.296 | HLA-A*11:01,HLA-A*30:01,HLA-A*03:01 | RBD |
| Spike | 372 | 9 | ASFSTFKCY | 7 | 0.480 | HLA-A*11:01,HLA-A*01:01HLA-B*46:01,HLA-B*15:02,HLA-B*15:01,HLA-B*35:01HLA-C*12:03 | RBD |
| Spike | 374 | 9 | FSTFKCYGV | 3 | 0.105 | HLA-A*02:06HLA-C*15:02,HLA-C*12:03 | RBD |
| Spike | 378 | 9 | KCYGVSPTK | 1 | 0.054 | HLA-A*30:01 | RBD |
| Spike | 379 | 9 | CYGVSPTKL | 3 | 0.350 | HLA-A*24:02HLA-C*07:02,HLA-C*14:02 | RBD |
| Spike | 392 | 9 | FTNVYADSF | 9 | 0.615 | HLA-A*26:01HLA-B*58:01,HLA-B*46:01,HLA-B*15:01,HLA-B*15:02HLA-C*07:02,HLA-C*03:04,HLA-C*03:03,HLA-C*12:03 | RBD |
| Spike | 394 | 9 | NVYADSFVI | 6 | 0.274 | HLA-A*02:06,HLA-A*32:01HLA-B*51:01HLA-C*15:02,HLA-C*03:04,HLA-C*12:03 | RBD |
| Spike | 399 | 9 | SFVIRGDEV | 1 | 0.042 | HLA-C*14:02 | RBD |
| Spike | 402 | 9 | IRGDEVRQI | 2 | 0.241 | HLA-C*07:02,HLA-C*06:02 | RBD |
| Spike | 409 | 9 | QIAPGQTGK | 2 | 0.241 | HLA-A*11:01,HLA-A*03:01 | RBD |
| Spike | 410 | 9 | IAPGQTGKI | 1 | 0.056 | HLA-B*51:01 | RBD |
| Spike | 411 | 9 | APGQTGKIA | 1 | 0.022 | HLA-B*07:02 | RBD |
| Spike | 413 | 9 | GQTGKIADY | 1 | 0.049 | HLA-B*15:01 | RBD |

| | | | | | | | |
|---|---|---|---|---|---|---|---|
| Spike | 417 | 9 | KIADYNYKL | 7 | 0.518 | HLA-A*02:01,HLA-A*02:07,HLA-A*02:06,HLA-A*32:01HLA-C*07:02,HLA-C*15:02,HLA-C*04:01 | RBD |
| Spike | 424 | 9 | KLPDDFTGC | 2 | 0.137 | HLA-A*02:07,HLA-A*02:06 | RBD |
| Spike | 425 | 9 | LPDDFTGCV | 3 | 0.098 | HLA-B*51:01,HLA-B*54:01,HLA-B*35:03 | RBD |
| Spike | 433 | 9 | VIAWNSNNL | 2 | 0.171 | HLA-C*03:04,HLA-C*03:03 | RBD |
| Spike | 444 | 9 | KVGGNYNYL | 2 | 0.088 | HLA-A*30:01HLA-C*15:02 | RBD |
| Spike | 448 | 9 | NYNYLYRLF | 4 | 0.439 | HLA-A*24:02HLA-C*07:02,HLA-C*14:02,HLA-C*06:02 | RBD |
| Spike | 453 | 9 | YRLFRKSNL | 4 | 0.301 | HLA-B*39:01HLA-C*07:02,HLA-C*14:02,HLA-C*06:02 | RBD |
| Spike | 454 | 9 | RLFRKSNLK | 3 | 0.296 | HLA-A*11:01,HLA-A*30:01,HLA-A*03:01 | RBD |
| Spike | 456 | 9 | FRKSNLKPF | 2 | 0.241 | HLA-C*07:02,HLA-C*06:02 | RBD |
| Spike | 458 | 9 | KSNLKPFER | 1 | 0.211 | HLA-A*11:01 | RBD |
| Spike | 462 | 9 | KPFERDIST | 2 | 0.052 | HLA-B*54:01,HLA-B*07:02 | RBD |
| Spike | 464 | 9 | FERDISTEI | 3 | 0.179 | HLA-B*40:01,HLA-B*40:02HLA-C*04:01 | RBD |
| Spike | 478 | 9 | TPCNGVEGF | 1 | 0.029 | HLA-B*35:01 | RBD |
| Spike | 481 | 9 | NGVEGFNCY | 2 | 0.064 | HLA-B*35:01,HLA-B*15:02 | RBD |
| Spike | 487 | 9 | NCYFPLQSY | 2 | 0.064 | HLA-B*35:01,HLA-B*15:02 | RBD |
| Spike | 489 | 9 | YFPLQSYGF | 4 | 0.409 | HLA-A*24:02HLA-C*07:02,HLA-C*14:02,HLA-C*04:01 | RBD |
| Spike | 490 | 9 | FPLQSYGFQ | 1 | 0.030 | HLA-B*54:01 | RBD |
| Spike | 495 | 9 | YGFQPTNGV | 5 | 0.324 | HLA-B*51:01HLA-C*03:04,HLA-C*06:02,HLA-C*04:01,HLA-C*12:03 | RBD |
| Spike | 497 | 9 | FQPTNGVGY | 3 | 0.187 | HLA-B*46:01,HLA-B*15:01,HLA-B*15:02 | RBD |
| Spike | 503 | 9 | VGYQPYRVV | 4 | 0.264 | HLA-B*51:01HLA-C*03:04,HLA-C*06:02,HLA-C*12:03 | RBD |
| Spike | 504 | 9 | GYQPYRVVV | 3 | 0.350 | HLA-A*24:02HLA-C*07:02,HLA-C*14:02 | RBD |
| Spike | 505 | 9 | YQPYRVVVL | 8 | 0.593 | HLA-A*02:07HLA-B*39:01HLA-C*07:02,HLA-C*03:04,HLA-C*04:01,HLA-C*03:03,HLA-C*06:02,HLA-C*12:03 | RBD |

| | | | | | | | |
|---|---|---|---|---|---|---|---|
| Spike | 507 | 9 | PYRVVVLSF | 1 | 0.156 | HLA-A*24:02 | RBD |
| Spike | 509 | 9 | RVVVLSFEL | 7 | 0.413 | HLA-A*02:07,HLA-A*02:06,HLA-A*32:01HLA-B*58:01HLA-C*15:02,HLA-C*03:03,HLA-C*03:04 | RBD |
| Spike | 511 | 9 | VVLSFELLH | 1 | 0.211 | HLA-A*11:01 | RBD |
| Spike | 512 | 9 | VLSFELLHA | 2 | 0.158 | HLA-A*02:03,HLA-A*02:01 | RBD |
| Spike | 515 | 9 | FELLHAPAT | 1 | 0.019 | HLA-B*40:02 | |
| Spike | 526 | 9 | GPKKSTNLV | 1 | 0.022 | HLA-B*07:02 | |
| Spike | 529 | 9 | KSTNLVKNK | 3 | 0.296 | HLA-A*11:01,HLA-A*30:01,HLA-A*03:01 | |
| Spike | 533 | 9 | LVKNKCVNF | 2 | 0.085 | HLA-B*15:01,HLA-B*15:02 | |
| Spike | 535 | 9 | KNKCVNFNF | 1 | 0.013 | HLA-A*32:01 | |
| Spike | 550 | 9 | GVLTESNKK | 2 | 0.241 | HLA-A*11:01,HLA-A*03:01 | |
| Spike | 554 | 9 | ESNKKFLPF | 3 | 0.092 | HLA-A*26:01HLA-B*35:01,HLA-B*15:02 | |
| Spike | 560 | 9 | LPFQQFGRD | 1 | 0.030 | HLA-B*54:01 | |
| Spike | 568 | 9 | DIADTTDAV | 1 | 0.028 | HLA-A*26:01 | |
| Spike | 576 | 9 | VRDPQTLEI | 4 | 0.318 | HLA-B*39:01HLA-C*07:02,HLA-C*06:02,HLA-C*04:01 | |
| Spike | 582 | 9 | LEILDITPC | 1 | 0.019 | HLA-B*40:02 | |
| Spike | 584 | 9 | ILDITPCSF | 5 | 0.229 | HLA-A*01:01HLA-B*35:01,HLA-B*15:02HLA-C*03:03,HLA-C*04:01 | |
| Spike | 587 | 9 | ITPCSFGGV | 2 | 0.137 | HLA-A*02:07,HLA-A*02:06 | |
| Spike | 590 | 9 | CSFGGVSVI | 5 | 0.268 | HLA-B*58:01,HLA-B*51:01HLA-C*15:02,HLA-C*03:04,HLA-C*12:03 | |
| Spike | 603 | 9 | NTSNQVAVL | 3 | 0.190 | HLA-C*03:04,HLA-C*03:03,HLA-C*12:03 | |
| Spike | 604 | 9 | TSNQVAVLY | 8 | 0.501 | HLA-A*11:01,HLA-A*01:01HLA-B*58:01,HLA-B*15:02,HLA-B*46:01,HLA-B*35:01,HLA-B*57:01HLA-C*12:03 | |

| | | | | | | |
|---|---|---|---|---|---|---|
| Spike | 612 | 9 | YQDVNCTEV | 6 | 0.409 | HLA-A*02:07,HLA-A*02:06,HLA-A*02:01HLA-B*39:01HLA-C*03:03,HLA-C*04:01 |
| Spike | 624 | 9 | IHADQLTPT | 1 | 0.018 | HLA-B*39:01 |
| Spike | 625 | 9 | HADQLTPTW | 4 | 0.158 | HLA-B*58:01,HLA-B*35:01,HLA-B*57:01HLA-C*04:01 |
| Spike | 628 | 9 | QLTPTWRVY | 3 | 0.113 | HLA-B*35:01,HLA-B*15:01,HLA-B*15:02 |
| Spike | 630 | 9 | TPTWRVYST | 2 | 0.052 | HLA-B*54:01,HLA-B*07:02 |
| Spike | 634 | 9 | RVYSTGSNV | 8 | 0.415 | HLA-A*02:03,HLA-A*30:01,HLA-A*32:01HLA-C*15:02,HLA-C*03:04,HLA-C*03:03,HLA-C*06:02,HLA-C*12:03 |
| Spike | 635 | 9 | VYSTGSNVF | 3 | 0.350 | HLA-A*24:02HLA-C*07:02,HLA-C*14:02 |
| Spike | 642 | 9 | VFQTRAGCL | 2 | 0.102 | HLA-C*14:02,HLA-C*04:01 |
| Spike | 643 | 9 | FQTRAGCLI | 3 | 0.169 | HLA-A*02:06HLA-B*39:01HLA-C*03:04 |
| Spike | 644 | 9 | QTRAGCLIG | 1 | 0.054 | HLA-A*30:01 |
| Spike | 652 | 9 | GAEHVNNSY | 2 | 0.063 | HLA-A*01:01HLA-B*35:01 |
| Spike | 654 | 9 | EHVNNSYEC | 1 | 0.018 | HLA-B*39:01 |
| Spike | 658 | 9 | NSYECDIPI | 3 | 0.190 | HLA-B*51:01HLA-C*15:02,HLA-C*03:04 |
| Spike | 660 | 9 | YECDIPIGA | 1 | 0.019 | HLA-B*40:02 |
| Spike | 664 | 9 | IPIGAGICA | 4 | 0.092 | HLA-B*35:01,HLA-B*07:02,HLA-B*54:01,HLA-B*35:03 |
| Spike | 666 | 9 | IGAGICASY | 5 | 0.235 | HLA-B*46:01,HLA-B*15:02,HLA-B*15:01,HLA-B*35:01HLA-C*12:03 |
| Spike | 679 | 9 | NSPRRARSV | 1 | 0.089 | HLA-C*06:02 |
| Spike | 680 | 9 | SPRRARSVA | 2 | 0.052 | HLA-B*54:01,HLA-B*07:02 |
| Spike | 683 | 9 | RARSVASQS | 1 | 0.054 | HLA-A*30:01 |
| Spike | 684 | 9 | ARSVASQSI | 3 | 0.259 | HLA-B*39:01HLA-C*07:02,HLA-C*06:02 |
| Spike | 685 | 9 | RSVASQSII | 4 | 0.158 | HLA-A*30:01HLA-B*58:01,HLA-B*57:01HLA-C*15:02 |

| | | | | | | |
|---|---|---|---|---|---|---|
| Spike | 687 | 9 | VASQSIIAY | 8 | 0.399 | HLA-A*01:01HLA-B*15:01,HLA-B*58:01,HLA-B*15:02,HLA-B*46:01,HLA-B*35:01HLA-C*03:03,HLA-C*12:03 |
| Spike | 689 | 9 | SQSIIAYTM | 4 | 0.174 | HLA-B*15:01,HLA-B*39:01,HLA-B*15:02HLA-C*03:03 |
| Spike | 691 | 9 | SIIAYTMSL | 15 | 0.909 | HLA-A*02:01,HLA-A*02:07,HLA-A*02:06,HLA-A*32:01HLA-B*15:01,HLA-B*35:03,HLA-B*15:02,HLA-B*46:01,HLA-B*07:02,HLA-B*39:01HLA-C*07:02,HLA-C*03:04,HLA-C*15:02,HLA-C*14:02,HLA-C*03:03 |
| Spike | 693 | 9 | IAYTMSLGA | 3 | 0.149 | HLA-B*54:01HLA-C*03:04,HLA-C*12:03 |
| Spike | 697 | 9 | MSLGAENSV | 5 | 0.261 | HLA-A*02:06HLA-B*51:01HLA-C*15:02,HLA-C*03:04,HLA-C*12:03 |
| Spike | 699 | 9 | LGAENSVAY | 6 | 0.306 | HLA-B*46:01,HLA-B*15:02,HLA-B*15:01,HLA-B*35:01HLA-C*03:03,HLA-C*12:03 |
| Spike | 704 | 9 | SVAYSNNSI | 4 | 0.140 | HLA-A*32:01HLA-B*07:02HLA-C*15:02,HLA-C*03:03 |
| Spike | 705 | 9 | VAYSNNSIA | 3 | 0.149 | HLA-B*54:01HLA-C*03:04,HLA-C*12:03 |
| Spike | 706 | 9 | AYSNNSIAI | 3 | 0.350 | HLA-A*24:02HLA-C*07:02,HLA-C*14:02 |
| Spike | 710 | 9 | NSIAIPTNF | 8 | 0.426 | HLA-B*58:01,HLA-B*15:02,HLA-B*46:01,HLA-B*35:01,HLA-B*57:01HLA-C*03:04,HLA-C*03:03,HLA-C*12:03 |
| Spike | 712 | 9 | IAIPTNFTI | 12 | 0.556 | HLA-A*02:06,HLA-A*32:01HLA-B*35:03,HLA-B*58:01,HLA-B*46:01,HLA-B*35:01,HLA-B*51:01,HLA-B*57:01HLA-C*15:02,HLA-C*03:03,HLA-C*03:04,HLA-C*12:03 |
| Spike | 714 | 9 | IPTNFTISV | 5 | 0.148 | HLA-B*35:03,HLA-B*54:01,HLA-B*07:02,HLA-B*35:01,HLA-B*51:01 |

| | | | | | | |
|---|---|---|---|---|---|---|
| Spike | 718 | 9 | FTISVTTEI | 11 | 0.737 | HLA-A*02:01,HLA-A*02:06,HLA-A*26:01HLA-B*46:01,HLA-B*58:01HLA-C*15:02,HLA-C*03:04,HLA-C*04:01,HLA-C*03:03,HLA-C*06:02,HLA-C*12:03 |
| Spike | 721 | 9 | SVTTEILPV | 6 | 0.348 | HLA-A*02:03,HLA-A*02:07,HLA-A*02:06,HLA-A*02:01HLA-C*15:02,HLA-C*12:03 |
| Spike | 725 | 9 | EILPVSMTK | 1 | 0.211 | HLA-A*11:01 |
| Spike | 727 | 9 | LPVSMTKTS | 1 | 0.030 | HLA-B*54:01 |
| Spike | 732 | 9 | TKTSVDCTM | 1 | 0.018 | HLA-B*39:01 |
| Spike | 733 | 9 | KTSVDCTMY | 3 | 0.104 | HLA-A*01:01HLA-B*58:01,HLA-B*57:01 |
| Spike | 734 | 9 | TSVDCTMYI | 5 | 0.253 | HLA-A*02:06HLA-B*58:01HLA-C*15:02,HLA-C*06:02,HLA-C*12:03 |
| Spike | 746 | 9 | STECSNLLL | 2 | 0.068 | HLA-A*01:01HLA-C*15:02 |
| Spike | 751 | 9 | NLLLQYGSF | 1 | 0.035 | HLA-B*15:02 |
| Spike | 755 | 9 | QYGSFCTQL | 3 | 0.350 | HLA-A*24:02HLA-C*07:02,HLA-C*14:02 |
| Spike | 757 | 9 | GSFCTQLNR | 1 | 0.211 | HLA-A*11:01 |
| Spike | 759 | 9 | FCTQLNRAL | 3 | 0.182 | HLA-B*35:03HLA-C*03:04,HLA-C*03:03 |
| Spike | 762 | 9 | QLNRALTGI | 1 | 0.035 | HLA-A*02:03 |
| Spike | 764 | 9 | NRALTGIAV | 3 | 0.259 | HLA-B*39:01HLA-C*07:02,HLA-C*06:02 |
| Spike | 773 | 9 | EQDKNTQEV | 1 | 0.018 | HLA-B*39:01 |
| Spike | 777 | 9 | NTQEVFAQV | 2 | 0.086 | HLA-A*02:06HLA-C*15:02 |
| Spike | 780 | 9 | EVFAQVKQI | 2 | 0.084 | HLA-A*26:01HLA-B*51:01 |
| Spike | 781 | 9 | VFAQVKQIY | 2 | 0.194 | HLA-C*07:02,HLA-C*14:02 |
| Spike | 782 | 9 | FAQVKQIYK | 3 | 0.296 | HLA-A*11:01,HLA-A*30:01,HLA-A*03:01 |
| Spike | 786 | 9 | KQIYKTPPI | 4 | 0.237 | HLA-A*02:01,HLA-A*02:06,HLA-A*32:01HLA-B*15:01 |
| Spike | 787 | 9 | QIYKTPPIK | 3 | 0.296 | HLA-A*11:01,HLA-A*30:01,HLA-A*03:01 |
| Spike | 789 | 9 | YKTPPIKDF | 2 | 0.241 | HLA-C*07:02,HLA-C*06:02 |

| | | | | | | |
|---|---|---|---|---|---|---|
| Spike | 794 | 9 | IKDFGGFNF | 1 | 0.060 | HLA-C*04:01 |
| Spike | 797 | 9 | FGGFNFSQI | 2 | 0.156 | HLA-B*51:01HLA-C*03:04 |
| Spike | 798 | 9 | GGFNFSQIL | 1 | 0.019 | HLA-C*12:03 |
| Spike | 803 | 9 | SQILPDPSK | 1 | 0.211 | HLA-A*11:01 |
| Spike | 810 | 9 | SKPSKRSFI | 1 | 0.089 | HLA-C*06:02 |
| Spike | 814 | 9 | KRSFIEDLL | 3 | 0.259 | HLA-B*39:01HLA-C*07:02,HLA-C*06:02 |
| Spike | 815 | 9 | RSFIEDLLF | 6 | 0.185 | HLA-A*32:01HLA-B*58:01,HLA-B*15:01,HLA-B*57:01HLA-C*15:02,HLA-C*12:03 |
| Spike | 817 | 9 | FIEDLLFNK | 1 | 0.211 | HLA-A*11:01 |
| Spike | 818 | 9 | IEDLLFNKV | 2 | 0.119 | HLA-B*40:01,HLA-B*40:02 |
| Spike | 821 | 9 | LLFNKVTLA | 4 | 0.295 | HLA-A*02:03,HLA-A*02:07,HLA-A*02:06,HLA-A*02:01 |
| Spike | 825 | 9 | KVTLADAGF | 2 | 0.024 | HLA-A*32:01HLA-B*57:01 |
| Spike | 826 | 9 | VTLADAGFI | 1 | 0.034 | HLA-C*15:02 |
| Spike | 827 | 9 | TLADAGFIK | 2 | 0.241 | HLA-A*11:01,HLA-A*03:01 |
| Spike | 829 | 9 | ADAGFIKQY | 1 | 0.026 | HLA-B*44:03 |
| Spike | 833 | 9 | FIKQYGDCL | 2 | 0.171 | HLA-C*03:04,HLA-C*03:03 |
| Spike | 852 | 9 | AQKFNGLTV | 2 | 0.104 | HLA-A*30:01HLA-B*15:01 |
| Spike | 853 | 9 | QKFNGLTVL | 4 | 0.278 | HLA-B*39:01HLA-C*07:02,HLA-C*06:02,HLA-C*12:03 |
| Spike | 857 | 9 | GLTVLPPLL | 2 | 0.208 | HLA-A*02:07,HLA-A*02:01 |
| Spike | 861 | 9 | LPPLLTDEM | 4 | 0.118 | HLA-B*35:01,HLA-B*51:01,HLA-B*35:03,HLA-B*07:02 |
| Spike | 865 | 9 | LTDEMIAQY | 5 | 0.177 | HLA-A*01:01HLA-B*35:01,HLA-B*15:02HLA-C*04:01,HLA-C*12:03 |
| Spike | 868 | 9 | EMIAQYTSA | 1 | 0.028 | HLA-A*26:01 |
| Spike | 869 | 9 | MIAQYTSAL | 17 | 0.941 | HLA-A*32:01,HLA-A*02:03,HLA-A*02:07,HLA-A*02:06,HLA-A*02:01HLA-B*15:01,HLA-B*35:03,HLA-B*15:02,HLA-B*46:01,HLA-B*07:02,HLA-B*39:01HLA- |

| | | | | | | |
|---|---|---|---|---|---|---|
| | | | | | | C*15:02,HLA-C*03:04,HLA-C*04:01,HLA-C*14:02,HLA-C*03:03,HLA-C*06:02 |
| Spike | 870 | 9 | IAQYTSALL | 7 | 0.373 | HLA-B*46:01,HLA-B*35:03,HLA-B*15:02HLA-C*15:02,HLA-C*03:03,HLA-C*03:04,HLA-C*12:03 |
| Spike | 871 | 9 | AQYTSALLA | 1 | 0.052 | HLA-A*02:06 |
| Spike | 874 | 9 | TSALLAGTI | 2 | 0.090 | HLA-B*51:01HLA-C*15:02 |
| Spike | 878 | 9 | LAGTITSGW | 2 | 0.070 | HLA-B*58:01,HLA-B*57:01 |
| Spike | 880 | 9 | GTITSGWTF | 6 | 0.323 | HLA-A*24:02,HLA-A*32:01HLA-B*58:01,HLA-B*15:01,HLA-B*15:02,HLA-B*57:01 |
| Spike | 886 | 9 | WTFGAGAAL | 12 | 0.531 | HLA-A*02:06HLA-B*15:01,HLA-B*15:02,HLA-B*07:02,HLA-B*35:01,HLA-B*39:01HLA-C*15:02,HLA-C*03:04,HLA-C*04:01,HLA-C*14:02,HLA-C*03:03,HLA-C*12:03 |
| Spike | 888 | 9 | FGAGAALQI | 5 | 0.336 | HLA-B*51:01HLA-C*03:04,HLA-C*03:03,HLA-C*06:02,HLA-C*12:03 |
| Spike | 890 | 9 | AGAALQIPF | 2 | 0.151 | HLA-B*46:01,HLA-B*15:01 |
| Spike | 892 | 9 | AALQIPFAM | 9 | 0.460 | HLA-B*35:03,HLA-B*58:01,HLA-B*15:02,HLA-B*46:01,HLA-B*35:01HLA-C*15:02,HLA-C*03:04,HLA-C*03:03,HLA-C*12:03 |
| Spike | 894 | 9 | LQIPFAMQM | 11 | 0.721 | HLA-A*02:06HLA-B*46:01,HLA-B*15:01,HLA-B*39:01,HLA-B*15:02HLA-C*07:02,HLA-C*15:02,HLA-C*03:04,HLA-C*03:03,HLA-C*06:02,HLA-C*12:03 |
| Spike | 896 | 9 | IPFAMQMAY | 6 | 0.233 | HLA-B*35:03,HLA-B*15:02,HLA-B*54:01,HLA-B*35:01,HLA-B*51:01HLA-C*03:03 |

| | | | | | | |
|---|---|---|---|---|---|---|
| Spike | 898 | 9 | FAMQMAYRF | 16 | 0.902 | HLA-A*24:02,HLA-A*32:01HLA-B*15:01,HLA-B*35:03,HLA-B*58:01,HLA-B*15:02,HLA-B*46:01,HLA-B*35:01,HLA-B*51:01,HLA-B*57:01HLA-C*03:04,HLA-C*04:01,HLA-C*14:02,HLA-C*03:03,HLA-C*06:02,HLA-C*12:03 |
| Spike | 901 | 9 | QMAYRFNGI | 2 | 0.048 | HLA-A*02:03,HLA-A*32:01 |
| Spike | 903 | 9 | AYRFNGIGV | 2 | 0.097 | HLA-A*30:01HLA-C*14:02 |
| Spike | 904 | 9 | YRFNGIGVT | 2 | 0.169 | HLA-B*39:01HLA-C*07:02 |
| Spike | 909 | 9 | IGVTQNVLY | 1 | 0.029 | HLA-B*35:01 |
| Spike | 915 | 9 | VLYENQKLI | 3 | 0.209 | HLA-A*02:03,HLA-A*02:07HLA-C*06:02 |
| Spike | 919 | 9 | NQKLIANQF | 1 | 0.035 | HLA-B*15:02 |
| Spike | 922 | 9 | LIANQFNSA | 3 | 0.117 | HLA-A*02:03,HLA-A*02:06HLA-B*54:01 |
| Spike | 923 | 9 | IANQFNSAI | 8 | 0.429 | HLA-B*46:01,HLA-B*51:01,HLA-B*35:03,HLA-B*15:02HLA-C*15:02,HLA-C*03:03,HLA-C*03:04,HLA-C*12:03 |
| Spike | 925 | 9 | NQFNSAIGK | 2 | 0.241 | HLA-A*11:01,HLA-A*03:01 |
| Spike | 937 | 9 | SLSSTASAL | 6 | 0.312 | HLA-A*02:03HLA-B*15:01,HLA-B*15:02,HLA-B*07:02HLA-C*03:04,HLA-C*03:03 |
| Spike | 939 | 9 | SSTASALGK | 3 | 0.296 | HLA-A*11:01,HLA-A*30:01,HLA-A*03:01 |
| Spike | 940 | 9 | STASALGKL | 4 | 0.224 | HLA-C*15:02,HLA-C*03:03,HLA-C*03:04,HLA-C*12:03 |
| Spike | 943 | 9 | SALGKLQDV | 5 | 0.276 | HLA-A*02:06HLA-C*15:02,HLA-C*03:03,HLA-C*03:04,HLA-C*12:03 |
| Spike | 950 | 9 | DVVNQNAQA | 1 | 0.028 | HLA-A*26:01 |
| Spike | 951 | 9 | VVNQNAQAL | 5 | 0.237 | HLA-B*35:03,HLA-B*07:02HLA-C*15:02,HLA-C*03:04,HLA-C*03:03 |
| Spike | 955 | 9 | NAQALNTLV | 1 | 0.056 | HLA-B*51:01 |

| | | | | | | |
|---|---|---|---|---|---|---|
| Spike | 956 | 9 | AQALNTLVK | 2 | 0.241 | HLA-A*11:01,HLA-A*03:01 |
| Spike | 958 | 9 | ALNTLVKQL | 1 | 0.035 | HLA-A*02:03 |
| Spike | 962 | 9 | LVKQLSSNF | 4 | 0.215 | HLA-B*46:01,HLA-B*15:02,HLA-B*15:01,HLA-B*35:01 |
| Spike | 964 | 9 | KQLSSNFGA | 1 | 0.052 | HLA-A*02:06 |
| Spike | 965 | 9 | QLSSNFGAI | 1 | 0.035 | HLA-A*02:03 |
| Spike | 968 | 9 | SNFGAISSV | 2 | 0.053 | HLA-C*15:02,HLA-C*12:03 |
| Spike | 969 | 9 | NFGAISSVL | 3 | 0.258 | HLA-A*24:02HLA-C*14:02,HLA-C*04:01 |
| Spike | 972 | 9 | AISSVLNDI | 1 | 0.035 | HLA-A*02:03 |
| Spike | 973 | 9 | ISSVLNDIL | 5 | 0.274 | HLA-B*58:01,HLA-B*57:01HLA-C*15:02,HLA-C*03:03,HLA-C*03:04 |
| Spike | 975 | 9 | SVLNDILSR | 1 | 0.211 | HLA-A*11:01 |
| Spike | 976 | 9 | VLNDILSRL | 6 | 0.457 | HLA-A*02:03,HLA-A*02:07,HLA-A*02:06,HLA-A*02:01HLA-B*46:01HLA-C*04:01 |
| Spike | 981 | 9 | LSRLDKVEA | 1 | 0.054 | HLA-A*30:01 |
| Spike | 983 | 9 | RLDKVEAEV | 3 | 0.268 | HLA-A*02:07,HLA-A*02:01HLA-C*04:01 |
| Spike | 989 | 9 | AEVQIDRLI | 3 | 0.145 | HLA-B*44:03,HLA-B*40:01,HLA-B*40:02 |
| Spike | 996 | 9 | LITGRLQSL | 3 | 0.190 | HLA-C*03:04,HLA-C*03:03,HLA-C*12:03 |
| Spike | 999 | 9 | GRLQSLQTY | 2 | 0.241 | HLA-C*07:02,HLA-C*06:02 |
| Spike | 1000 | 9 | RLQSLQTYV | 4 | 0.295 | HLA-A*02:03,HLA-A*02:07,HLA-A*02:06,HLA-A*02:01 |
| Spike | 1004 | 9 | LQTYVTQQL | 5 | 0.337 | HLA-B*40:01,HLA-B*15:01,HLA-B*39:01HLA-C*03:04,HLA-C*03:03 |
| Spike | 1005 | 9 | QTYVTQQLI | 6 | 0.268 | HLA-B*58:01,HLA-B*51:01,HLA-B*57:01HLA-C*15:02,HLA-C*06:02,HLA-C*12:03 |
| Spike | 1012 | 9 | LIRAAEIRA | 1 | 0.054 | HLA-A*30:01 |
| Spike | 1014 | 9 | RAAEIRASA | 3 | 0.156 | HLA-A*30:01HLA-B*54:01HLA-C*03:03 |
| Spike | 1016 | 9 | AEIRASANL | 3 | 0.145 | HLA-B*44:03,HLA-B*40:01,HLA-B*40:02 |

| | | | | | | |
|---|---|---|---|---|---|---|
| Spike | 1018 | 9 | IRASANLAA | 1 | 0.018 | HLA-B*39:01 |
| Spike | 1020 | 9 | ASANLAATK | 3 | 0.296 | HLA-A*11:01,HLA-A*30:01,HLA-A*03:01 |
| Spike | 1021 | 9 | SANLAATKM | 8 | 0.401 | HLA-B*46:01,HLA-B*15:02,HLA-B*35:03,HLA-B*35:01HLA-C*15:02,HLA-C*03:03,HLA-C*03:04,HLA-C*12:03 |
| Spike | 1026 | 9 | ATKMSECVL | 1 | 0.054 | HLA-A*30:01 |
| Spike | 1048 | 9 | HLMSFPQSA | 2 | 0.158 | HLA-A*02:03,HLA-A*02:01 |
| Spike | 1050 | 9 | MSFPQSAPH | 2 | 0.131 | HLA-B*46:01,HLA-B*35:01 |
| Spike | 1052 | 9 | FPQSAPHGV | 6 | 0.208 | HLA-B*35:03,HLA-B*54:01,HLA-B*07:02,HLA-B*35:01,HLA-B*51:01HLA-C*04:01 |
| Spike | 1054 | 9 | QSAPHGVVF | 11 | 0.639 | HLA-A*32:01HLA-B*15:01,HLA-B*58:01,HLA-B*15:02,HLA-B*46:01,HLA-B*35:01,HLA-B*57:01HLA-C*07:02,HLA-C*03:04,HLA-C*03:03,HLA-C*12:03 |
| Spike | 1055 | 9 | SAPHGVVFL | 5 | 0.401 | HLA-C*07:02,HLA-C*03:04,HLA-C*04:01,HLA-C*03:03,HLA-C*12:03 |
| Spike | 1056 | 9 | APHGVVFLH | 1 | 0.029 | HLA-B*35:01 |
| Spike | 1059 | 9 | GVVFLHVTY | 3 | 0.097 | HLA-A*32:01HLA-B*15:01,HLA-B*15:02 |
| Spike | 1060 | 9 | VVFLHVTYV | 8 | 0.468 | HLA-A*02:03,HLA-A*02:07,HLA-A*02:06,HLA-A*02:01HLA-B*54:01HLA-C*15:02,HLA-C*06:02,HLA-C*12:03 |
| Spike | 1062 | 9 | FLHVTYVPA | 5 | 0.325 | HLA-A*02:03,HLA-A*02:07,HLA-A*02:06,HLA-A*02:01HLA-B*54:01 |
| Spike | 1065 | 9 | VTYVPAQEK | 3 | 0.296 | HLA-A*11:01,HLA-A*30:01,HLA-A*03:01 |
| Spike | 1073 | 9 | KNFTTAPAI | 1 | 0.013 | HLA-A*32:01 |
| Spike | 1086 | 9 | KAHFPREGV | 2 | 0.088 | HLA-A*30:01HLA-C*15:02 |
| Spike | 1087 | 9 | AHFPREGVF | 3 | 0.259 | HLA-B*39:01HLA-C*07:02,HLA-C*06:02 |

| | | | | | | |
|---|---|---|---|---|---|---|
| Spike | 1088 | 9 | HFPREGVFV | 1 | 0.042 | HLA-C*14:02 |
| Spike | 1089 | 9 | FPREGVFVS | 2 | 0.059 | HLA-B*35:01,HLA-B*54:01 |
| Spike | 1094 | 9 | VFVSNGTHW | 1 | 0.156 | HLA-A*24:02 |
| Spike | 1095 | 9 | FVSNGTHWF | 12 | 0.890 | HLA-A*24:02,HLA-A*26:01HLA-B*46:01,HLA-B*35:01,HLA-B*15:01,HLA-B*15:02HLA-C*07:02,HLA-C*03:04,HLA-C*04:01,HLA-C*03:03,HLA-C*06:02,HLA-C*12:03 |
| Spike | 1096 | 9 | VSNGTHWFV | 2 | 0.092 | HLA-B*58:01HLA-C*15:02 |
| Spike | 1099 | 9 | GTHWFVTQR | 1 | 0.211 | HLA-A*11:01 |
| Spike | 1101 | 9 | HWFVTQRNF | 3 | 0.350 | HLA-A*24:02HLA-C*07:02,HLA-C*14:02 |
| Spike | 1102 | 9 | WFVTQRNFY | 1 | 0.042 | HLA-C*14:02 |
| Spike | 1106 | 9 | QRNFYEPQI | 1 | 0.089 | HLA-C*06:02 |
| Spike | 1109 | 9 | FYEPQIITT | 1 | 0.152 | HLA-C*07:02 |
| Spike | 1113 | 9 | QIITTDNTF | 5 | 0.243 | HLA-A*26:01HLA-B*46:01,HLA-B*15:02,HLA-B*15:01,HLA-B*35:01 |
| Spike | 1120 | 9 | TFVSGNCDV | 1 | 0.042 | HLA-C*14:02 |
| Spike | 1121 | 9 | FVSGNCDVV | 8 | 0.519 | HLA-A*02:03,HLA-A*02:07,HLA-A*02:06,HLA-A*02:01HLA-C*15:02,HLA-C*03:03,HLA-C*03:04,HLA-C*12:03 |
| Spike | 1130 | 9 | IGIVNNTVY | 3 | 0.166 | HLA-B*46:01,HLA-B*15:02,HLA-B*35:01 |
| Spike | 1137 | 9 | VYDPLQPEL | 3 | 0.254 | HLA-C*07:02,HLA-C*14:02,HLA-C*04:01 |
| Spike | 1147 | 9 | SFKEELDKY | 1 | 0.042 | HLA-C*14:02 |
| Spike | 1158 | 9 | NHTSPDVDL | 1 | 0.018 | HLA-B*39:01 |
| Spike | 1161 | 9 | SPDVDLGDI | 1 | 0.011 | HLA-B*35:03 |
| Spike | 1169 | 9 | ISGINASVV | 1 | 0.034 | HLA-C*15:02 |
| Spike | 1171 | 9 | GINASVVNI | 2 | 0.048 | HLA-A*02:03,HLA-A*32:01 |

| | | | | | | |
|---|---|---|---|---|---|---|
| Spike | 1173 | 9 | NASVVNIQK | 1 | 0.211 | HLA-A*11:01 |
| Spike | 1175 | 9 | SVVNIQKEI | 1 | 0.034 | HLA-C*15:02 |
| Spike | 1181 | 9 | KEIDRLNEV | 4 | 0.197 | HLA-A*02:06HLA-B*44:03,HLA-B*40:01,HLA-B*40:02 |
| Spike | 1185 | 9 | RLNEVAKNL | 2 | 0.048 | HLA-A*02:03,HLA-A*32:01 |
| Spike | 1188 | 9 | EVAKNLNES | 1 | 0.028 | HLA-A*26:01 |
| Spike | 1189 | 9 | VAKNLNESL | 2 | 0.171 | HLA-C*03:04,HLA-C*03:03 |
| Spike | 1192 | 9 | NLNESLIDL | 2 | 0.158 | HLA-A*02:03,HLA-A*02:01 |
| Spike | 1201 | 9 | QELGKYEQY | 1 | 0.026 | HLA-B*44:03 |
| Spike | 1206 | 9 | YEQYIKWPW | 2 | 0.046 | HLA-B*44:03,HLA-B*40:02 |
| Spike | 1207 | 9 | EQYIKWPWY | 1 | 0.035 | HLA-B*15:02 |
| Spike | 1208 | 9 | QYIKWPWYI | 3 | 0.396 | HLA-A*24:02HLA-C*07:02,HLA-C*06:02 |
| Spike | 1209 | 9 | YIKWPWYIW | 3 | 0.179 | HLA-A*24:02,HLA-A*32:01HLA-B*57:01 |
| Spike | 1212 | 9 | WPWYIWLGF | 4 | 0.092 | HLA-B*35:01,HLA-B*54:01,HLA-B*35:03,HLA-B*07:02 |
| Spike | 1213 | 9 | PWYIWLGFI | 1 | 0.156 | HLA-A*24:02 |
| Spike | 1216 | 9 | IWLGFIAGL | 2 | 0.307 | HLA-A*24:02HLA-C*07:02 |
| Spike | 1217 | 9 | WLGFIAGLI | 1 | 0.035 | HLA-A*02:03 |
| Spike | 1218 | 9 | LGFIAGLIA | 1 | 0.030 | HLA-B*54:01 |
| Spike | 1219 | 9 | GFIAGLIAI | 1 | 0.042 | HLA-C*14:02 |
| Spike | 1220 | 9 | FIAGLIAIV | 8 | 0.550 | HLA-A*02:03,HLA-A*02:07,HLA-A*02:06,HLA-A*02:01HLA-B*46:01HLA-C*15:02,HLA-C*03:04,HLA-C*12:03 |
| Spike | 1221 | 9 | IAGLIAIVM | 4 | 0.210 | HLA-B*35:01,HLA-B*35:03HLA-C*03:04,HLA-C*03:03 |
| Spike | 1223 | 9 | GLIAIVMVT | 1 | 0.123 | HLA-A*02:01 |
| Spike | 1224 | 9 | LIAIVMVTI | 1 | 0.013 | HLA-A*32:01 |

| | | | | | | |
|---|---|---|---|---|---|---|
| Spike | 1225 | 9 | IAIVMVTIM | 8 | 0.426 | HLA-B*35:03,HLA-B*58:01,HLA-B*15:02,HLA-B*46:01,HLA-B*35:01HLA-C*03:04,HLA-C*03:03,HLA-C*12:03 |
| Spike | 1229 | 9 | MVTIMLCCM | 2 | 0.047 | HLA-B*35:03,HLA-B*15:02 |
| Spike | 1236 | 9 | CMTSCCSCL | 1 | 0.085 | HLA-A*02:07 |
| Spike | 1237 | 9 | MTSCCSCLK | 2 | 0.266 | HLA-A*11:01,HLA-A*30:01 |
| Spike | 1248 | 9 | CSCGSCCKF | 2 | 0.070 | HLA-B*58:01,HLA-B*57:01 |
| Spike | 1257 | 9 | DEDDSEPVL | 1 | 0.100 | HLA-B*40:01 |
| Spike | 1262 | 9 | EPVLKGVKL | 2 | 0.033 | HLA-B*35:03,HLA-B*07:02 |
| Spike | 1264 | 9 | VLKGVKLHY | 2 | 0.085 | HLA-B*15:01,HLA-B*15:02 |

Supplementary Table 3. Information of HLA types

| HLA_Type | iNeo_Epi_Count | netMHCPan_Epi_Count | Shared_Epi_Count | Union_Epi_Count |
|---|---|---|---|---|
| HLA-A*11:01 | 59 | 57 | 51 | 65 |
| HLA-A*24:02 | 81 | 64 | 49 | 96 |
| HLA-C*07:02 | 185 | 116 | 89 | 212 |
| HLA-A*02:01 | 99 | 51 | 48 | 102 |
| HLA-B*46:01 | 97 | 107 | 58 | 146 |
| HLA-C*03:04 | 214 | 113 | 88 | 239 |
| HLA-B*40:01 | 14 | 27 | 13 | 28 |
| HLA-C*06:02 | 161 | 93 | 71 | 183 |
| HLA-A*02:07 | 134 | 65 | 46 | 153 |
| HLA-C*03:03 | 198 | 113 | 85 | 226 |
| HLA-C*04:01 | 95 | 108 | 47 | 156 |
| HLA-B*58:01 | 60 | 52 | 45 | 67 |
| HLA-B*51:01 | 67 | 61 | 39 | 89 |
| HLA-A*30:01 | 116 | 67 | 56 | 127 |
| HLA-A*02:06 | 141 | 71 | 63 | 149 |
| HLA-B*15:01 | 113 | 65 | 63 | 115 |
| HLA-C*14:02 | 65 | 123 | 51 | 137 |
| HLA-B*15:02 | 372 | 87 | 86 | 373 |
| HLA-A*02:03 | 85 | 55 | 44 | 96 |
| HLA-A*01:01 | 27 | 56 | 21 | 62 |
| HLA-C*15:02 | 165 | 107 | 79 | 193 |
| HLA-B*54:01 | 84 | 52 | 38 | 98 |
| HLA-A*03:01 | 41 | 40 | 26 | 55 |
| HLA-B*35:01 | 123 | 94 | 78 | 139 |

| | | | | |
|---|---|---|---|---|
| HLA-A*26:01 | 30 | 70 | 20 | 80 |
| HLA-B*44:03 | 9 | 25 | 8 | 26 |
| HLA-B*48:01 | 0 | 55 | 0 | 55 |
| HLA-B*07:02 | 52 | 32 | 28 | 56 |
| HLA-B*40:02 | 34 | 29 | 17 | 46 |
| HLA-C*12:03 | 158 | 138 | 98 | 198 |
| HLA-B*39:01 | 60 | 62 | 39 | 83 |
| HLA-A*32:01 | 107 | 95 | 60 | 142 |
| HLA-B*35:03 | 61 | 78 | 36 | 103 |
| HLA-B*57:01 | 98 | 40 | 34 | 104 |
